# Raw milk kefir: microbiota, bioactive peptides, and immune modulation

**DOI:** 10.1101/2022.10.19.512868

**Authors:** Ton Baars, Betty van Esch, Luuk van Ooijen, Zuomin Zhang, Pieter Dekker, Sjef Boeren, Mara Diks, Johan Garssen, Kasper Hettinga, Remco Kort

**Author notes:** Shared first authors (Ton Baars, Betty van Esch).

## Abstract

This study aims to characterize the microbial and peptidomic composition of raw milk kefir, and to address the potential anti-allergic effects of raw milk kefir using validated research models for food allergy. Raw milk kefir was produced after incubation with a defined freeze-dried starter culture. Kefir was sampled during fermentation at seven-time intervals. For comparison, kefir was also prepared from boiled milk (100 °C; 5 sec). Peptide compositions were determined for the raw and heated milk, plus kefir end products made from these milks. In a murine food allergy model, the two kefir end products were investigated on their allergy modulating effects. In both kefirs, we identified amplicon sequence variants identical to those in the starter, matching the bacteria *Lactococcus lactis, Streptococcus thermophilus, Leuconostoc* and the yeast *Debaromyces*. In raw milk kefir, additional sequence variants of *Lactococcus lactis* and the yeasts *Pichia* and *Galactomyces* could be identified, which were absent in heated milk kefir. Analysis of peptide compositions in kefirs indicated that the number and intensity of peptides drastically increased after fermentation. Heating of milk negatively affected the diversity of the peptide composition in kefir. Only raw milk kefir suppressed the acute allergic skin response to the food allergen ovalbumin in sensitised mice. These effects coincided with differences in the T-cell compartment, with lower percentages of activated Th1 cells and IFNg production after treatment with kefir made from heated milk. The results of this study indicate specific properties of raw milk kefir that may contribute to its additional health benefits.

## 1. Introduction

Fermentation is the oldest way of milk conservation, transforming milk into foods, like kefir and yoghurt, as well as fresh or ripened cheese. Defined starter cultures (SC) based on selected microorganisms were not used until the early 1900s, implicating that fermentation was initiated by microorganisms originating from the environment and equipment (1). Over the last decades, there has been an increased interest for kefir, a drinkable milk product that originates from Eastern Europe, Turkey, or the Caucasus region. In contrast to yoghurt fermentation, which is based on thermophilic bacteria, kefir is fermented by mesophilic bacteria as well as yeasts. Wild starters of kefir are based on kefir grains or SCOBY, a symbiotic community of bacteria and yeast containing a matrix of the exopolysaccharide kefiran and proteins. The microbial species in these kefir grains include lactic acid bacteria, acetic acid bacteria and yeasts. Little kefir is still made this way, and the most common method to produce kefir on an industrial scale is based on fermentation with defined, often freeze-dried SC or through back-slopping, (2). These SC are based on a selection of species and strains, including *Streptococcus*, *Lactobacillus*, and *Lactococcus* as well as *Kluvyeromyces* and *Debaryomyces* (3). During the anaerobic fermentation process, there is multiplication of bacteria and yeast. According to the Codex Alimentarius (2003) kefir should contain at least 10^7^ cfu/mL of bacteria, and at least 10^4^ cfu/mL of yeast.

Fermented dairy belongs to the group of ‘functional foods’ or ‘nutraceuticals’ (4) The health impact of fermented food consumption is widely accepted, and benefits may be based on multiple mechanisms (5). For kefir, lactose fermenting micro-organisms are responsible for the transformation of lactose into lactate and other organic acids, causing a drop in pH. During fermentation, peptides are produced because of proteolytic degradation of mainly caseins (6), but also short chain fatty acids, ethanol, and vitamins are generated (4). Some kefir peptides show biological activity, including angiotensin-converting enzyme (ACE)-inhibitory, antimicrobial, immunomodulating, opioid, mineral binding, antioxidant, and antithrombotic effects (6,7). Overall health claims associated with kefir consumption were reviewed from different perspectives (8–12), but studies on the impact on food allergy are missing. Pre-clinical studies on kefir show promising results on infections, glucose and cholesterol pathway, intestinal microbiota composition, inflammatory, allergy related molecules, like IgE and immune modulation in the gut (8). In a study utilizing an ovalbumin sensitisation murine asthma model, it was found that mice receiving intra-gastric kefir had lower levels of airway hyper-responsiveness (AHR) than control mice, and, impressively, had lower levels of AHR than the positive control group receiving an anti-asthma drug (13). The impact of kefir on gastro-intestinal health is based on changes in the gut microbiome, the improvement of gut dysbiosis, and improved gut barrier (4). Altered gut colonization and gut dysbiosis are associated with increased risk of immune related diseases, like asthma and allergies (14). In healthy volunteers, two weeks of kefir consumption changed the level of several cytokines, indicating an increase of the Th1 and a decrease of the Th2 immune response (15). In young rats, kefir consumption improved the intestinal mucosal and systemic immune responses after ingestion of cholera toxin (16). In murine studies, kefir consumption reduced the asthma response due to its anti-inflammatory and anti-allergic effects (13,17). Although pre-clinical studies on kefir show immune modulation after kefir consumption as well as reduced inflammation and airway hyperresponsiveness in allergic asthma, it is currently unknown whether kefir has the potential to prevent food allergic symptoms. Kefir is usually made from processed milk, including a boiling or pasteurisation step, using skimmed milk, or homogenized full fat milk. Knowledge about the impact of kefir made from unheated and non-homogenized full fat milk is missing, and there is an increasing amount of literature on the impact of processing of milk on its composition and its relation to human health (18–20). Various epidemiological studies have shown reduced incidences of asthma and allergies in children who consumed bovine raw milk (21). After the raw milk was heated, the protection was lost, even in farm children (22). These epidemiologic observations were confirmed in recent studies where we showed anti-allergic effects of raw milk in murine food allergy models and a clinical pilot study with 11 allergic children (23–25). It could be shown that the protection is related to the whey fraction of the milk, containing a wide range of heat-sensitive proteins (24).

The aim of this study was to evaluate the impact of kefir based on SC, either made from raw or boiled whole milk. Thereto, the processing of raw milk kefir (RMK) and heated milk kefir (HMK) were characterized through the analysis of the kefir microbial and peptide composition. Both RMK and HMK were investigated on their allergy protective effect in a murine food allergy model. Microbial and peptide composition were used to hypothesize on potential mechanisms of allergy protection.

## 2. Materials and Methods

### 2.1 Kefir production and sampling

Kefir was produced at a commercial dairy plant, the Raw Milk Company (De Lutte, The Netherlands). The raw milk came from a mix of 80 dairy cows in one morning milking. Raw bulk milk was cooled at the farm to approximately 25 °C, and the time between the end of milking and the start of fermentation was less than two hours. The farm was organic certified, according to SKAL/EU regulations. The kefir produced this way is referred to as ‘Raw Milk Kefir’ (RMK). RMK was produced by a 2% addition to raw milk of a freeze-dried starter culture from the Christian Hansen company (Hoersholm, Denmark), eXact^®^ 2 kefir. Fermentation was done at 28 °C for 24 hours. Before bottling, the kefir was gently mixed. Bottles were stored chilled to 4 °C. The Hansen company declares in the product information (Version: 2 PI EU EN 03-03-2018) the presence of five different bacterial species: *Leuconostoc, Streptococcus thermophilus, Lactococcus lactis* subsp*. cremoris, L. lactis* subsp. *lactis, L. lactis* biovar*. Diacetylactis*, and the yeast *Debaryomyces hansenii*.

At seven different time points samples were taken both during the fermentation process of the raw milk as well as during the storage of the final kefir storage. To control the fermentation process, the pH was measured every 1 to 3 hours. To compare the impact of heating, the raw milk was heated (HM) to produce HM kefir (HMK). Milk used for HMK originated from the same batch of raw milk. After thermisation, 3L milk was shortly boiled and cooled back to 28 °C. In contrast to RMK, HMK was only sampled at two timepoints: milk after heating, before inoculation, and at the end of kefir processing after 24 hours. A schematic overview of the fermentation process and the samples collected and analysed in this study is displayed in Fig 1.

**Figure 1.**
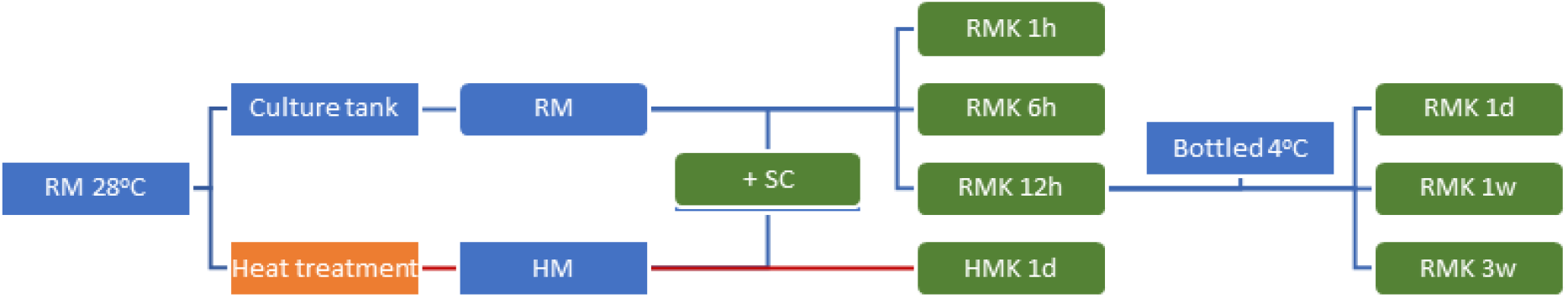
Timeline of the processing and sample collection steps for kefir production. Green boxes indicate the samples taken for microbial community analysis. Abbreviations: RM = raw milk; HM = heated milk; SC = defined kefir starter culture (Chr. Hansen); RMK = raw milk kefir (final product obtained after 1 day of fermentation); HMK = heated milk kefir (final product obtained after 1 day of fermentation); 1h, 1 hour; 1 d, 1 day; 1w/3w 1 or 3 weeks.

### 2.2 Enumeration of total bacteria, lactic acid bacteria and yeasts

Three different types of nutrient agar plates were used to determine the number of total bacteria, lactic acid bacteria and yeasts. Petri dishes with culture media and the NaCl solutions (0.9%, pH=7.0) for the dilution series were delivered by Biotrading Benelux BV (Mijdrecht, The Netherlands). Total bacterial counts were determined on Tryptic Soy Agar (TSA), lactic acid bacteria on de Man, Rogosa and Sharpe (MRS) agar, and yeasts on Sabouraud Dextrose Agar (SDA) agar with chloramphenicol to prevent bacterial growth. For each sample (Fig 1), serial dilutions were made and colony forming units were determined after 24 to 30 hours of incubation at 30 °C.

### 2.3 16S rRNA and ITS amplicon sequencing

#### DNA extraction, mock community, and genomic DNA spike-in

Samples were taken during milk fermentation and immediately transferred to dry ice in 2 mL tubes to stop further fermentation. At the end of the day all samples were stored at −18 °C. After two weeks the whole set of samples was transported on dry ice to the laboratory and stored at −80 °C before further analysis. The microbiota composition was determined by amplicon sequencing of the bacterial v3-v4 region from the 16S rRNA gene and the fungal internal transcribed spacer 1 (ITS). The DNA isolation efficiency, read depth and taxonomic assignment of the amplicons was determined by use of a mock microbial community (Microbial community standard II log distribution, zymoBIOMICS™, Irvine, CA, United States). A genomics spike-in (Microbial community DNA standard II log distribution, zymoBIOMICS™) was added to the samples as a control. Three replicates of three different HMK samples were determined to evaluate the reproducibility of bacterial microbiota and mycobiota compositions in this study.

A specific milk bacterial DNA isolation kit (Norgen Biotek, Thorold, Canada) was used with slight modifications of the manufacturer’s protocol. The following amount of sample was used. For unfermented milk 1 mL per sample, for fermented samples 0.2 mL with the addition of 0.8 mL phosphate buffer. For the defined SC, 20 mg was added to 1 mL phosphate buffered saline solution. The manufacturer recommendation of addition of 75 μL mock microbial community suspension was used, equivalent to a total of 220 ng of genomic DNA. All fermented samples were passed through a 21-gauge syringe (Sigma-Aldrich, St. Louis, MS, Unites States) to reduce the viscosity. After centrifuging for 2 min at 20,000 g the supernatant was removed. The cell pellet was resuspended in 100 μL resuspension solution A from the DNA isolation kit and transferred to a 2 mL screw cap tube preloaded with 0.5 g autoclaved glass beads. The samples were chilled on ice for 1 minute, followed by a 10 second bead shaking treatment with the speed set to 6 meters per second on a Fastprep®-24 5G Instrument (MP Biomedicals, Solon, OH, United States). Cycles of chilling and shaking were performed eight times. After mechanical cell lysis, the DNA isolation kit manufacture protocol was applied as specified for Gram-positive and unknown bacterial species.

#### Real-time polymerase chain reaction

The spike-in DNA derived reads per sample were aimed to be limited to 10% of the total reads per sample. Therefore, a quantitative SYBR based qPCR on the 16S rRNA gene was deployed on the gDNA from the samples and on the 5 times 10-fold serial diluted spike-in DNA. Per reaction, 1 μL gDNA template and 19 μL mastermix was used per reaction, which consists of 10 μL SYBR mastermix from SensiFast SYBR^®^ Hi-Rox-Kit (Meridian Bioscience, London, United Kingdom) 0.8 μL forward F357: “CCTACGGGAGGCAGCAG” (10 μM), 0.8 μL reverse R518 “ATTACCGCGGCTGCTGG” (10 μM) primer and 5 μL nucleotide-DNase free water. The qPCR was performed on a 7300 RT-PCR system (Applied Biosystems, Waltham, MA, United States) with the amplification setup of an initial start for 10 min at 95 °C followed by 40 amplification cycles with denaturation at 95 °C for 5s, annealing at 58 °C for 10s and elongation for 30s at 73 °C. The spike-in was added to all samples except for the RM and HM. A complete overview of the sample Ct values and estimated and measured concentration of added spike-in can be found in supplemental file 1.

#### DNA Quality control, library preparation and sequencing

DNA quality control, metagenome amplicons amplification, library preparation and sequencing were performed by Macrogen (Seoul, Korea). The double-stranded DNA binding dye, Picogreen (cat. #P7589, Invitrogen, Walthan, MA, United States) was added and DNA was quantified by Victor 3 fluorometry (Waltham, MA, United States). Amplicons of the V3-V4 region of the 16S rRNA gene and the and the ITS1 region and library prep were generated by Herculase II Fusion DNA Polymerase Nextera XT Index Kit V2 (Illumina, San Diego, CA, USA). The products of the library prep were validated by an Agilent Tapestation D1000 (Santa Clara, CA, USA). Paired-end sequencing was performed with 2 x 300 base pairs (bp) per sequence on the Illumina MiSeq System platform. The bcl2fastq package from Illumina was used to demultiplex the sequence data and to convert the base calls binary into FASTQ files.

#### Data processing

Demultiplexed and adapter trimmed sequences were visualized and loaded into Qiime2-2020.2 (26) through the paired-end manifest method. Based on the quality plots, sequences were truncated at the median nucleotide Phred score of <30. The trimmed sequences (removal of primers) were denoised, paired end joined, followed by removal of chimeras by the DADA2 plugin (27). Taxonomic annotation of the amplicons was performed by a Naive Bayes based model, trained on the 16SrRNA V3-V4 region extracted from the SILVA 138 database (28). For the ITS classification the model was trained on full length sequences from the dynamic clustered UNITE-V8 database (29). Taxonomic classification was performed on an Amazon Web Services (Seattle, WA, USA), AWS EC2 QIIME 2 Core – version 2020.2 (26). The feature table was exported to a biological observation matrix (.biom) file and converted into a tab separated file which was concatenated with the assigned taxonomy and amplicon sequence variant (ASV). The spike-in ASVs were identified and removed. The ASVs were blasted against the National Centre for Biotechnology Information (NCBI) database and the taxonomic assignment was manually curated. The relative abundance of ASV per sample were calculated by the ratio of reads ASV / total non-spike-in ASV reads per sample. Visualization of the data was carried out by (Graphpad Prism version: 6), (San Diego, CA, USA)).

#### Data availability

All raw sequence data generated in this project are available at NCBI under the bio-project number: PRJNA716278 https://www.ncbi.nlm.nih.gov/bioproject/PRJNA716278. The ASVs and meta-data are present in the supplemental file 1.

### 2.4 Kefir peptidomics

#### Sample preparation

The –80 °C stored samples were thawed and subsequently centrifuged at 16,000 g and 4 °C for 10 min to remove caseins and fat (7). Remaining proteins were precipitated by addition of an equal volume of 200 g/L trichloroacetic acid in milli-Q water, followed by centrifugation at 3,000 g and 4 °C for 10 min. The peptide fraction in the supernatant was cleaned-up with a solid phase extraction (SPE) on C18+ Stage tip columns (prepared in-house), (30). SPE was carried out as described before (31), using 100 μL supernatant of each sample. Lastly, peptides were reconstituted to 50 μL with 1 mL/L formic acid in water.

#### LC-MS/MS analysis

Peptide samples were analysed with nano LC-MS/MS as described before, with minor adjustments (32). In brief, 4 μL peptide solution was loaded onto a 0.10 × 250 mm ReproSil-Pur 120 C18-AQ 1.9 μm beads analytical column (prepared in-house) at 825 bar. Peptides were separated in a gradient changing from 9 to 34% acetonitrile in water with 0.1% formic acid over 50 min (Thermo nLC1000). Full scan FTMS spectra were obtained using a Q-Exactive HFX (Thermo electron, San Jose, CA, USA) in positive mode between 380 and 1,400 *m/z* at resolution 60,000. The 25 most abundant positively charged peaks (2-5+) in the MS scan were isolated and fragmented (HCD) with an isolation width of 1.2 *m/z* and 24% normalized collision energy. MSMS scans were recorded at resolution 15,000 in data-dependent mode with a threshold of 1.2 × 10^5^ and 15 s exclusion duration for the selected *m/z* +/− 10 ppm.

#### Data processing and analysis

Raw LC-MS/MS data files resulting from the peptidomics analysis were processed using MaxQuant v1.6.1.0 (33). For an initial search, a database comprising all *Bos taurus* protein sequences from the UniProtKB (34) was used. For this search, minimum peptide length was set to 8 and maximum peptide length to 25 amino acids.

In a second search, a new database was created comprising only the protein sequences of which peptides were identified in the first, initial search. This new database comprised 39 protein sequences and was used for a new search in which the minimum peptide length was set to 4 and the maximum peptide length to 45 amino acids. The rationale behind this search strategy was to comprehensively map the complete kefir peptidome with short processing times. In both searches, oxidation of methionine, n-terminal acetylation, deamidation of asparagine and glutamine and phosphorylation of serine and threonine were set as variable modifications. Peptides were filtered on identification score (> 80) and only peptides from proteins with > 5 unique peptides were retained.

Bioactive properties of identified peptide sequences were searched using the Milk Bioactive Peptide Database (MBPDB; http://mbpdb.nws.oregonstate.edu), (35). The search type used in the database was ‘precursor’ with a similarity threshold of 100%. Results were filtered on the requirement that the length of the bioactive sequence > 4 amino acids and that the length of the identified sequence < 2 × length bioactive sequence.

Sequence logos were created for the P1 and P1’ positions of the identified peptides. N- and C-terminals of the peptides were summed, and intensities of the amino acids were plotted using the R package ggseqlogo, version 0.1 (36).

### 2.5 Murine food allergy model

Three-week-old, specific pathogen-free, female C3H/HeOuJ mice (The Jackson Laboratory, Bar Harbor, ME, USA) were housed at the animal facility of Utrecht University (Utrecht, The Netherlands) in filter-topped makrolon cages (one cage/group, n = 6-8/cage) with standard chip bedding, Kleenex tissues and a plastic shelter, on a 12 h light/dark cycle with access to food (‘Rat and Mouse Breeder and Grower Expanded’; Special Diet Services, Witham, UK) and water ad libitum. Upon arrival, mice were randomly assigned to the 4 experimental groups and were habituated to the laboratory conditions for 6 days prior to the start of the study. All animal procedures were approved by the Ethical Committee for Animal Research of the Utrecht University and complied with the European Directive 2010/63/EU on the protection of animals used for scientific purposes (AVD108002015346).

On experimental days 0, 7, 14, 21 and 28, mice (n = 8/group) were orally sensitised, by using a blunt needle, to the hen’s egg protein ovalbumin (OVA) (20 mg/0.5 mL PBS; grade V; Sigma-Aldrich) using cholera toxin (CT; 15 μg/0.5 mL; List Biological Laboratories, Campbell, CA, United States) as an adjuvant. Sham-sensitised mice (PBS/PBS received CT alone (15 μg/0.5 mL PBS). The following experimental groups (treatment/sensitisation and challenge): PBS/PBS, PBS/OVA (allergic control), RMK/OVA and HMK/OVA. Milk and kefir samples were kept at –18 °C and thawed at the day of challenging. For the treatment with the different kefirs, PBS, RMK or HMK were given 3x/week by oral gavage from day −1 till 32. On day 33, mice were challenged i.d. in both ear pinnae with OVA (10 μg/20 μL PBS) to evaluate the acute allergic symptoms after i.d. challenge. Allergen-specific IgE in blood and percentages of activated Th1, Th2, Foxp3+ regulatory cells and IFNg production were measured on day 34, 16 hours after oral OVA challenge (50 mg OVA/0.5 mL PBS) in MLN or spleen. A schematic overview of the experimental timeline is shown in Fig 2.

**Figure 2.**
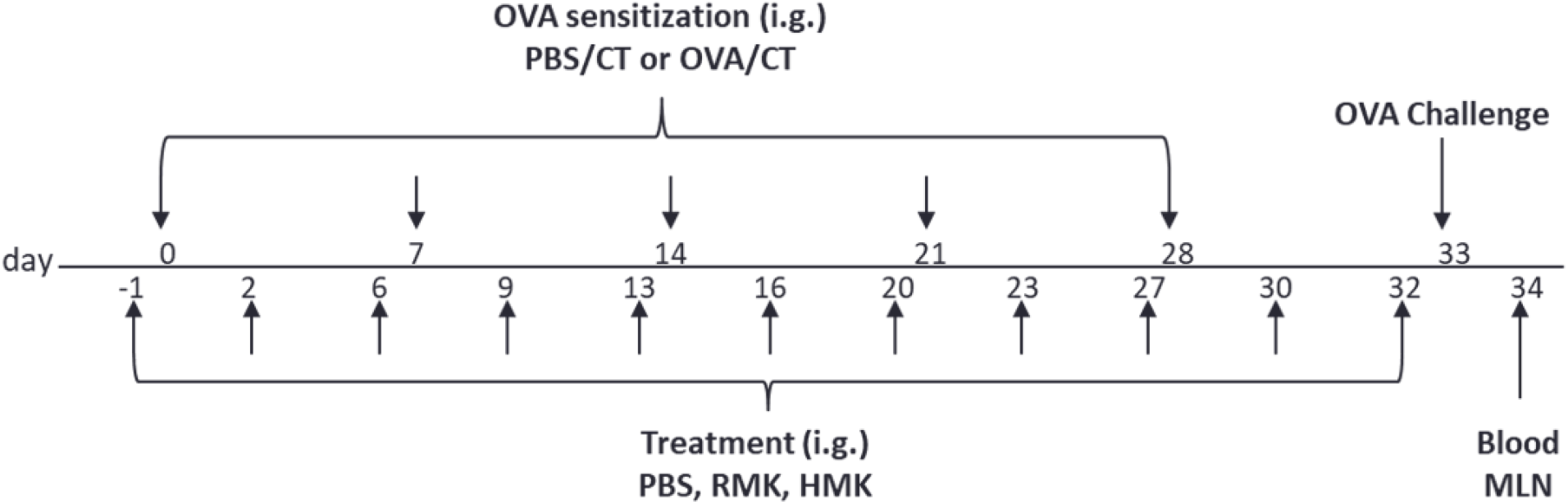
Schematic overview of the study design. Female C3H/HeOuJ mice were PBS treated or orally sensitised to OVA on day 0, 7, 14, 21, 28 using CT as adjuvant. The experimental set-up resulted in 4 groups: Sham-sensitised (PBS/PBS), OVA sensitised control mice (PBS/OVA), OVA sensitised RMK treated mice (RMK/OVA) and OVA sensitised HMK treated mice (HMK/OVA). PBS, RMK or HMK were given 3x/week by oral gavage (i.g.) from day −1 till 32. On day 33 mice were challenged i.d. with OVA to evaluate the allergic reaction. The acute allergic skin response, anaphylactic shock symptoms and body temperature are evaluated at 1 hour. Allergen-specific IgE, cytokine production, percentages of activated Th1 and Th2 cells were assessed at day 34, 16 hours after oral OVA challenge. OVA = ovalbumin; CT = cholera toxin. RMK = raw milk kefir; HMK = Heated milk Kefir; i.d. = intradermal; i.g. = intragastracilly by oral gavage.

#### Evaluation of the acute allergic response

To assess the severity of the acute allergic symptoms upon intradermal OVA challenge, both ear pinnae were injected with OVA and the acute allergic skin response, anaphylactic shock symptoms and body temperature were measured. The acute allergic skin response, expressed as Δ ear swelling (μm), was calculated by subtracting the mean basal ear thickness from the mean ear thickness measured 1 h after intradermal challenge. Ear thickness at both timepoints was measured in duplicate for each ear using a digital micrometer (Mitutoyo, Veenendaal, The Netherlands). To perform the intradermal challenge and both ear measurements, mice were anesthetized using inhalation of isoflurane (Abbott, Breda, The Netherlands). The severity of the anaphylactic shock symptoms was scored 30 min after the intradermal challenge by using a previously described, validated, scoring table (37).

#### Measurement of serum OVA-specific IgE

Blood collected via cheek puncture prior to sacrifice (day 34) was centrifuged at 10.000 rpm for 10 min. Serum was subsequently obtained and stored at −20 °C until analysis of OVA-specific IgE and IgG levels by means of ELISA as described elsewhere (25).

#### Th2 and Th1 percentages and *ex vivo* IFNg production

Single cell splenocyte and mesenteric lymph node (MLN) suspensions were obtained by passing the organs through a 70 μm filter. After red blood cell lysis, cells were blocked for 20 min in PBS containing 1% BSA and 5% FCS. 8×10^5^ Cells were plated per well and incubated for 30 min at 4 °C with different antibodies (eBioscience, Breda, The Netherlands or BD, Alphen aan de Rijn, The Netherlands, unless otherwise stated) against CD4, CD25, CD69, CXCR3, T1ST2, Foxp3 and isotype controls were used. Flow cytometry was performed using FACS Canto II (BD, Alphen aan den Rijn, The Netherlands) and analysed using FACSDiva software (BD). For cytokine release, splenocytes (8 × 10^5^ per well) were incubated in RPMI1640 supplemented with penicillin (100 U/mL), streptomycin (100 μg/mL), and 10% FBS. Cells were stimulated for 5 days with OVA (10 ug/mL). Supernatants were harvested for cytokine measurements by ELISA according to the manufacturer’s recommendations (eBioscience).

#### Statistical analysis

Experimental results are expressed as mean ± SEM, as individual data points or as box-and-whisker Tukey plots when data were not normally distributed. Differences between pre-selected groups were statistically determined using one-way ANOVA, followed by Bonferroni’s multiple comparisons test for preselected groups. OVA-specific IgE, OVA-specific IgG1 levels and anaphylactic shock scores were analysed using Kruskal-Wallis test for non-parametric data followed by Dunn’s multiple comparisons test for pre-selected groups. Statistical analyses were performed using GraphPad Prism software (version 7.03; GraphPad Software, San Diego, CA, USA) and results were considered statistically significant when *p* < 0.05.

## 3. Results

### 3.1 Microbial growth and acidification during fermentation and storage

The growth of lactic acid bacteria at 28 °C and concomitant production of lactic acid showed a typical pattern: a relatively slow reduction of the pH during the first hour, followed by a rapid decrease to pH 4.3 at 24 hours. A short lag phase was evident for yeast, but lactic acid bacteria show exponential growth from the first hour of incubation for starter cultures grown on lactose. The storage period from one day to three weeks at 4 °C was characterized by a further slow decline of the pH (Fig 3). The total increase in total bacterial load during the 24 hours of fermentation at 28 °C for RMK was about five orders of magnitude, from approximately 10^3^ to 10^8^ CFU/mL. An increase between four and five orders of magnitude in bacterial load can also be derived from the 16S rRNA gene-based cycle threshold values (Ct) of the quantitative PCR, as the Ct values decreased from 24 to 10 in the first 24 hours during the fermentation process. RMK and HMK do not significantly differ in Ct values or CFU counts of total or lactic acid bacteria after 1 day of fermentation. However, the HMK showed about 40-fold higher levels of yeasts (0.8 x 10^6^ CFU/mL) than RMK (0.2 x 10^5^ CFU/mL) after one day of fermentation at 28 °C. During storage of bottled RMK at 4 °C from 24 hours to three weeks a reduction of CFU count of lactic acid bacteria was observed, while the CFU count of yeasts further increased during three weeks of storage.

**Figure 3.**
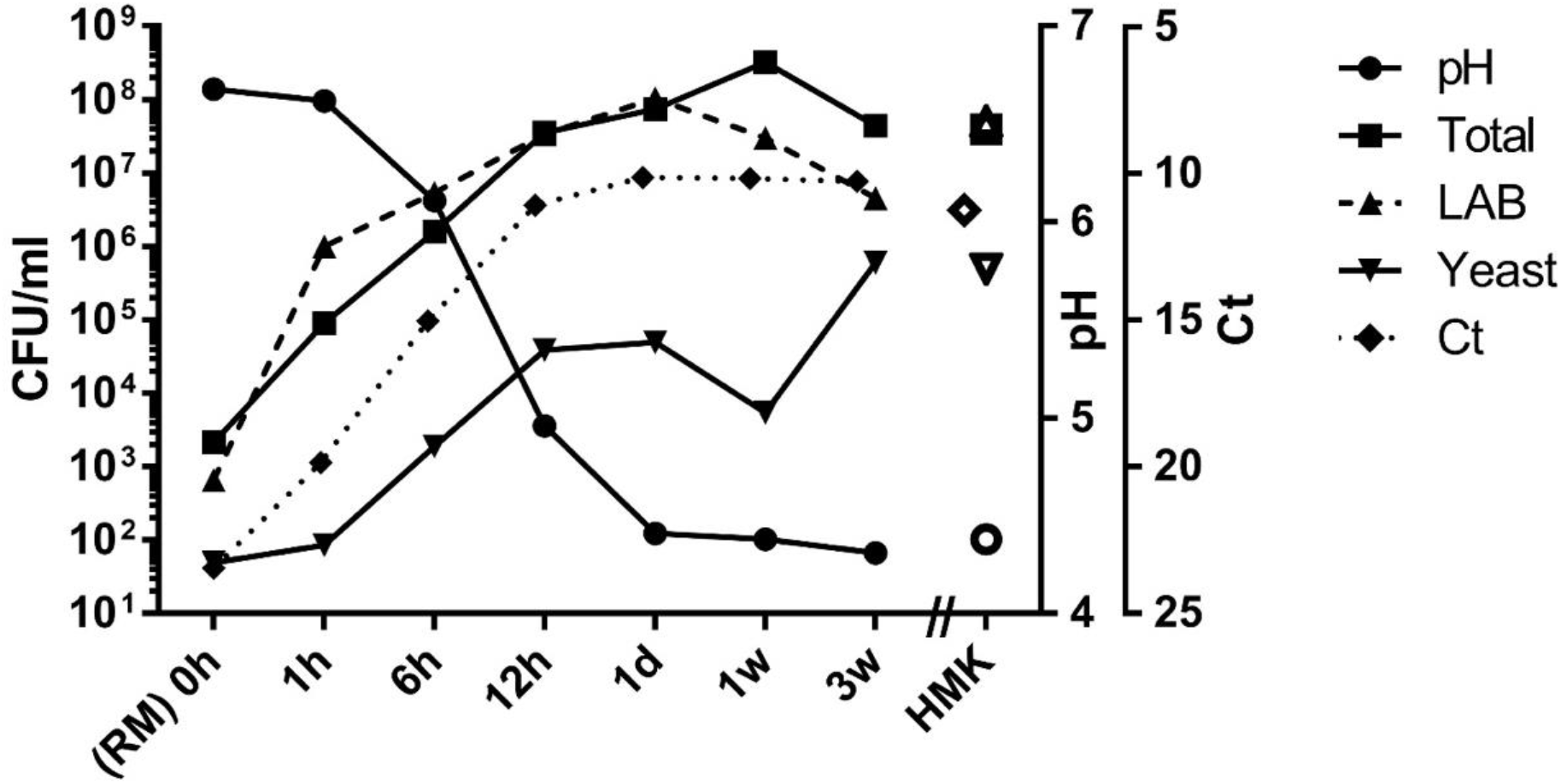
Acidity, microbial plate counts and cycle threshold values (Ct) during growth and storage of raw milk kefir. The acidity is expressed in pH-values (circles), the total microbial plate count (squares), LAB, lactic acid bacteria count (up-pointing triangles), and yeast plate counts (down-pointing triangles) in colony forming units per millilitre (CFU/mL), cycle threshold (Ct) values for the number of cycles required to exceed background for the qPCR of the bacterial 16S rRNA gene have been indicated on the inversed right y-axis (diamonds).The X-axis displays the time in hours after addition of the starter culture (SC). Raw milk (RM) is equivalent to the fermented product at time = 0 hours. The data for 0 to 24 hours (1 day) refer to raw milk fermentation at 28 °C, followed by storage at 4 °C from 1 day to three weeks. Data for heated milk kefir (HMK) obtained by 24 hours of fermentation at 28 °C have been indicated by open symbols.

### 3.2 Composition of microbiota during fermentation and storage of raw milk kefir

#### Drastic reduction of microbial species richness during fermentation

Analysis of the amplicon sequence variants (ASVs) of the starter culture (SC) led to the identification of eleven microbial ASVs, belonging to four bacterial species and one fungal ASV, including the bacteria *Streptococcus thermophilus, Lactococcus lactis, Lactococcus, Leuconostoc* and yeast *Debaromyce*s (Fig 4). These microbial species coincide with the five microbial species declared on the label of the used kefir starter culture. RM and HM both showed an extremely high richness of over 100 microbial species (supplemental file 1). However, it should be noted that the total viable count of bacteria and fungi in the RM was limited to values of only 10^3^ and 10^2^ CFU/mL, respectively (Fig 3). The process of kefir fermentation drastically reduced the microbial richness in the sample to 10 abundant microbial species, and after the first 12 hours of fermentation there was a limited change in composition, both in bacteria and in fungi (Fig 4; supplemental file 2). From this time point, the bacterial and fungal microbiota composition in the kefir is predominantly described by the ASVs that represent the five microbial species of the starter culture.

**Figure 4.**
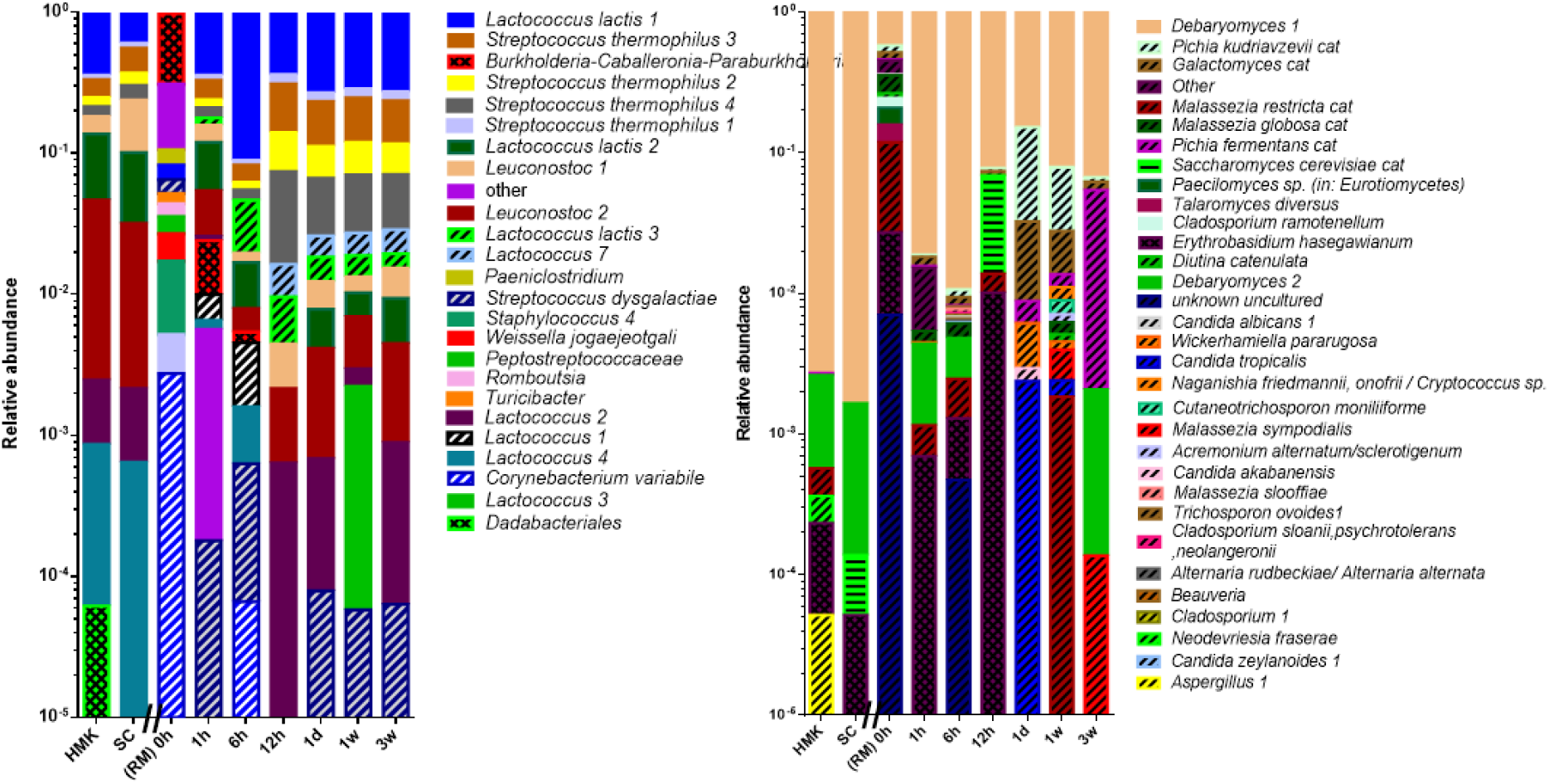
Logarithmic bar chart of the microbial composition during fermentation and storage of raw milk kefir. **A)** The relative number amplicon sequence variants (ASVs) determined by the bacterial V3-V4 region of the 16S rRNA gene amplicon, and **B)** the fungal ITS1 region. Genus and species names have been indicated in italics. The numbers behind the species names correspond to the number of observed unique ASVs. The label cat indicates that multiple ASVs have been concatenated to a single annotation. The Y-axis displays relative abundance in logarithmic order. The X-axis displays samples with time after addition of the starter culture. SC = defined starter culture. HMK = heated milk kefir after 24 hours of fermentation. Dashed lines indicate ASVs, which are not present in the starter culture and could originate from raw milk. Horizontal lines indicate ASVs that were assigned to species present in the spike-in. Chequered bars indicate microbial species that do not originate from RM.

#### Unique bacterial and fungal species in raw milk kefir

Further inspection of the data led to the identification of several bacterial and fungal species, which were present in RMK samples, but absent in HMK samples. Strikingly, all RMK samples from end-product at the first day to three weeks contained bacterial ASVs of *Lactococcus lactis* (ASV # 3; 0.6%)*, Lactococcus* (ASV # 7; 0.8%), both of which were absent in HMK. Although the relative abundances of the latter ASVs were low, the overall bacterial load has increased with five orders of magnitude to 10^8^ CFU/mL, leading to substantial presence of these bacteria in the final raw milk kefir products. In addition, ASVs from the yeasts *Pichia kudriavzevii* (12%) and *Galactomyces* (2.3%) could be identified in the final RMK products, but not in HMK (Fig 4). These two yeasts unique to raw milk kefir were represented by three and six distinct ASVs, respectively, while all the abundant bacterial and fungal ASVs identified in HMK were limited to those present in the starter culture. Minor amounts of the bacterium *Streptococcus dysgalactiae* were present in raw milk and RMK products as well the yeasts *Malessazia restricta* and *Malessazia globosa*. Finally, a few bacterial and fungal species could be identified as contaminants, including the bacterial genus *Burkholderia* and the yeast *Erythrobasidium hasegawianum*, which were present in milk with low overall microbial loads, but decreased during the process of fermentation and could not be identified in final products.

### 3.3 Composition of peptides in milk and kefir end products

#### Fermentation increases the number of peptides in kefir

Peptide analysis resulted in the identification of 2,525 different bovine peptides, of which 478 peptides were identified in all samples (Fig 5A). In addition, 1,039 peptides were only identified in kefir samples and 202 peptides were only found in milk, showing that fermentation of the milk results in a drastic increase in the number of unique peptides. This also holds for the total abundance of the peptides, which was several magnitudes higher in the kefir samples (Fig 5B). RM and RMK have a higher number of unique peptides when compared to HM and HMK, respectively (Fig 5A). Nevertheless, the total peptide intensity was higher in HMK than in RMK (Fig 5B).

**Figure 5.**
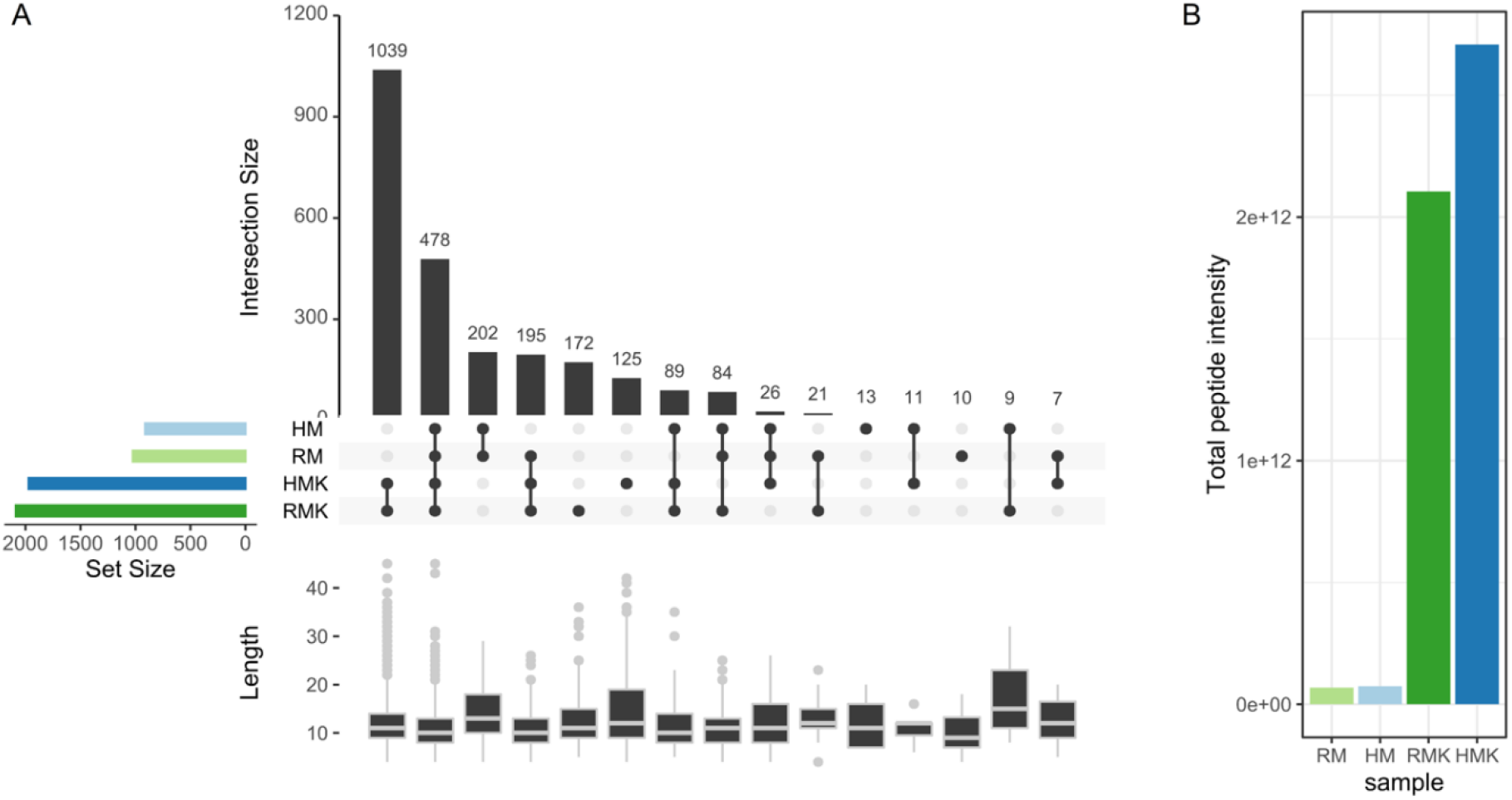
(A) UpSet plot visualizing the intersections of identified peptides in the different samples. Vertical bars represent the number of unique peptides identified in the intersection of samples shown underneath each bar. Horizontal bars represent the total number of peptides identified in each sample. Box plots underneath each section show the peptide lengths in the respective section. (B) Total peptide intensity per sample. With raw milk (RM), heated milk (HM), raw milk kefir (RMK), and heated milk kefir (HMK).

#### Most kefir peptides originate from caseins

Regarding the precursor proteins, it can be noted that most peptides originate from the caseins. This holds for both the number of peptides only identified in kefir (Table 1) and the peptide intensity (Fig 6 and supplemental file 3). Fermentation of the milk causes a notable increase in the number of unique peptides and the intensity of peptides from caseins, glycosylation-dependent cell adhesion molecule 1 (Glycam-1), osteopontin, and polymeric immunoglobulin receptor (PIGR).

**Figure 6.**
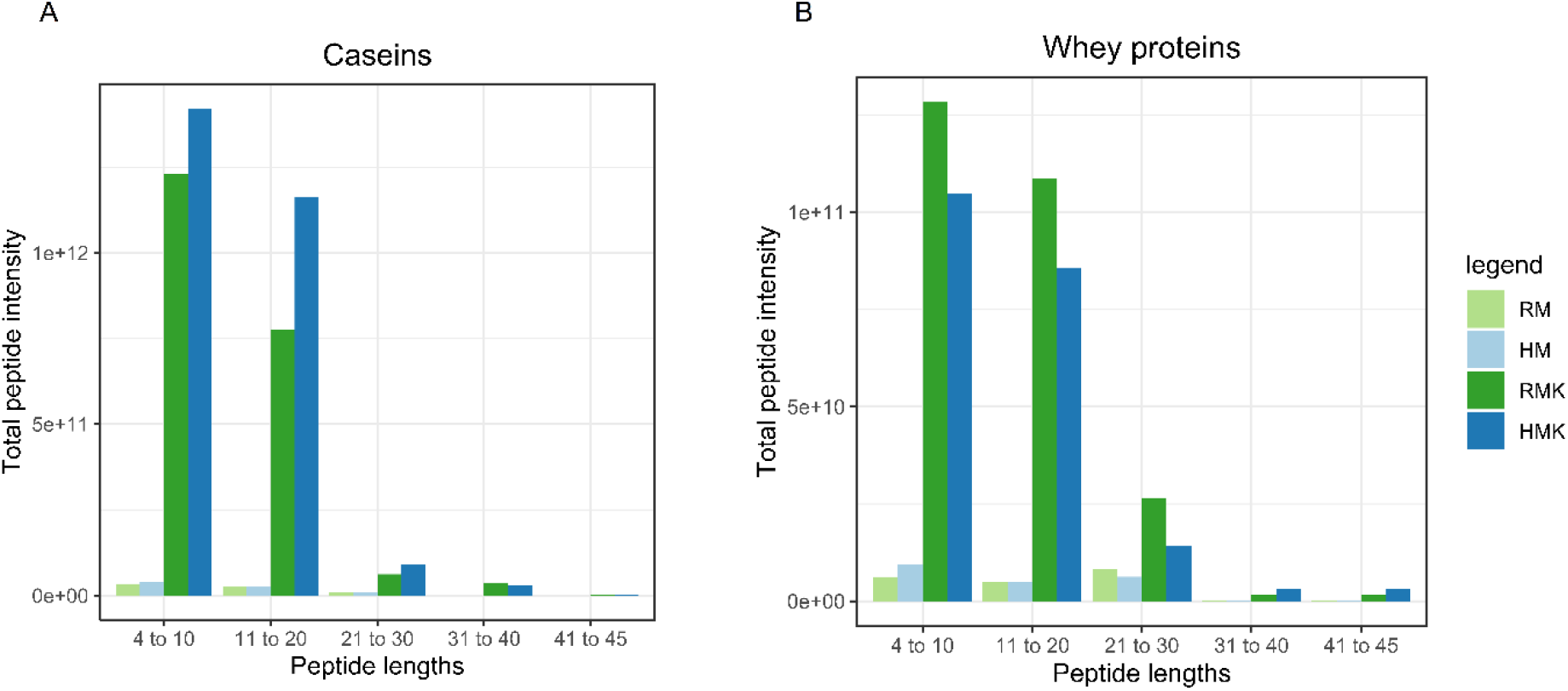
Bar plots of summed peptide intensity per binned range of peptide lengths. With peptides from (A) caseins and (B) whey proteins.

**Table 1.**
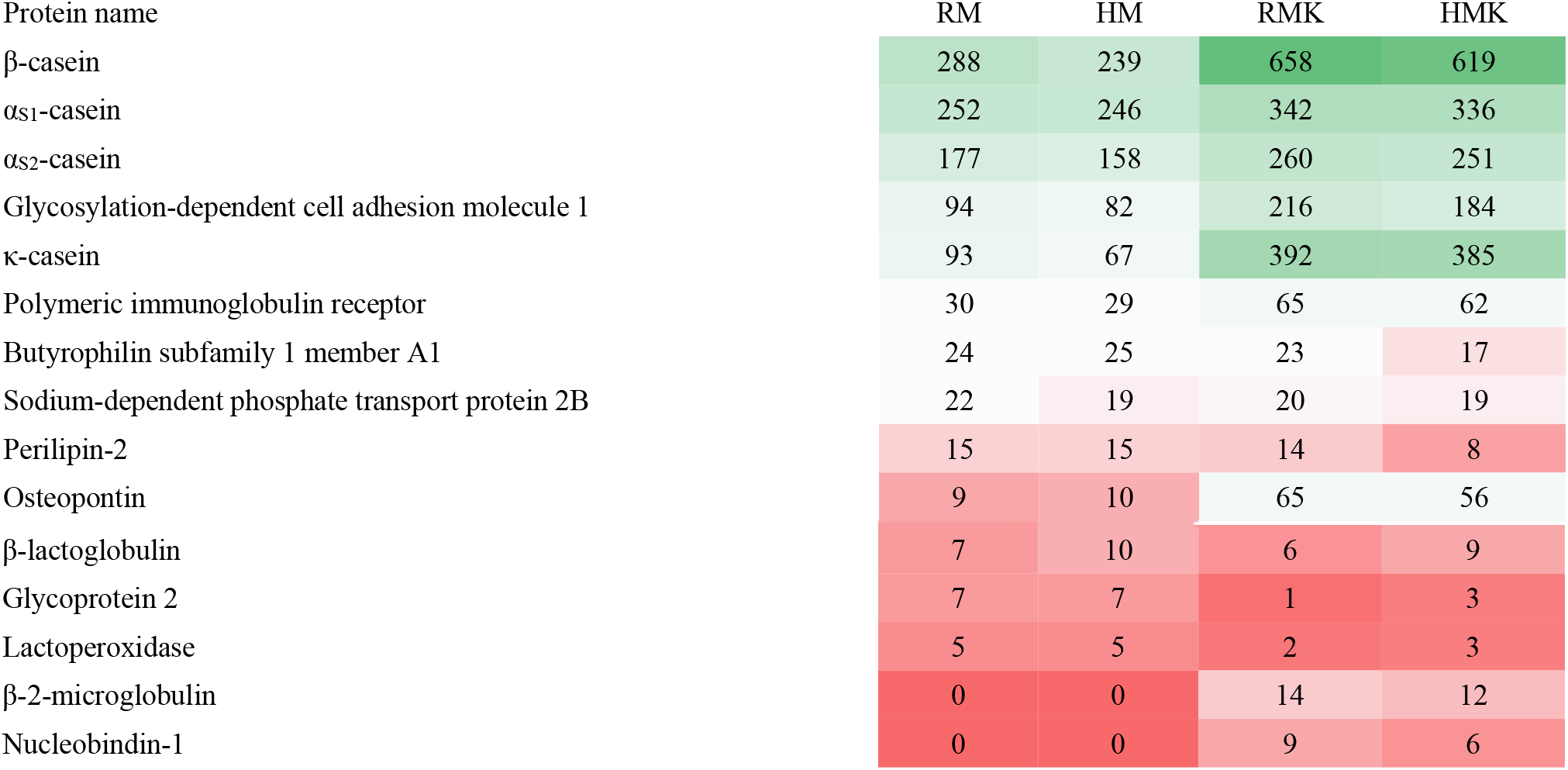
Precursor proteins of kefir peptides with the number of identified peptides in raw milk (RM), heated milk (HM), raw milk kefir (RMK), and heated milk kefir (HMK). Shown are proteins with at least 5 identified peptides in one of the 4 samples, arranged on number of identified peptides in RM. Colour gradient represents high numbers (green) to low numbers (red) of unique peptides.

#### Raw milk kefir shows a higher number of bioactive peptides

Whey peptides with a length up to 30 amino acids are more abundant in RMK than in HMK, whereas the opposite is true for peptides from caseins (Figure 6). Nevertheless, regarding peptides longer than 30 amino acids, peptides from whey proteins are more abundant in HMK than in RMK and again, the opposite is observed for peptides from caseins. Next, the presence of bioactive peptides was determined and their intensities in RMK and HMK compared (supplemental file 4). The most frequently found bioactivities were ACE-inhibitory, antioxidant, and antimicrobial activity. Comparing HMK with RMK, showed that HMK comprises a larger number of bioactive sequences that are more abundant in HMK than in RMK. Comparing bioactivities of peptides identified uniquely in either HMK or RMK, showed that RMK has a larger number of bioactive peptides.

### 3.4 Murine food allergy model

#### Treatment with RMK suppressed the acute allergic skin response

In mice treated with PBS, sensitisation with OVA (PBS/OVA) resulted in an acute allergic skin response (Fig 7A), and anaphylactic shock symptoms (Fig 7C) upon i.d. challenge compared to PBS/PBS mice. Sensitised mice treated with RMK (RMK/OVA) showed a reduced acute allergic skin response compared to PBS/OVA allergic mice. Although the anaphylactic shock score was lower in RMK/OVA mice, no significant effects were observed on anaphylactic shock symptoms and body temperature compared to PBS/OVA mice. Sensitised mice treated with HMK (HMK/OVA) showed no effect on the acute allergic skin response indicating that heating of milk before fermentation affected the protective effect of kefir.

**Figure 7.**
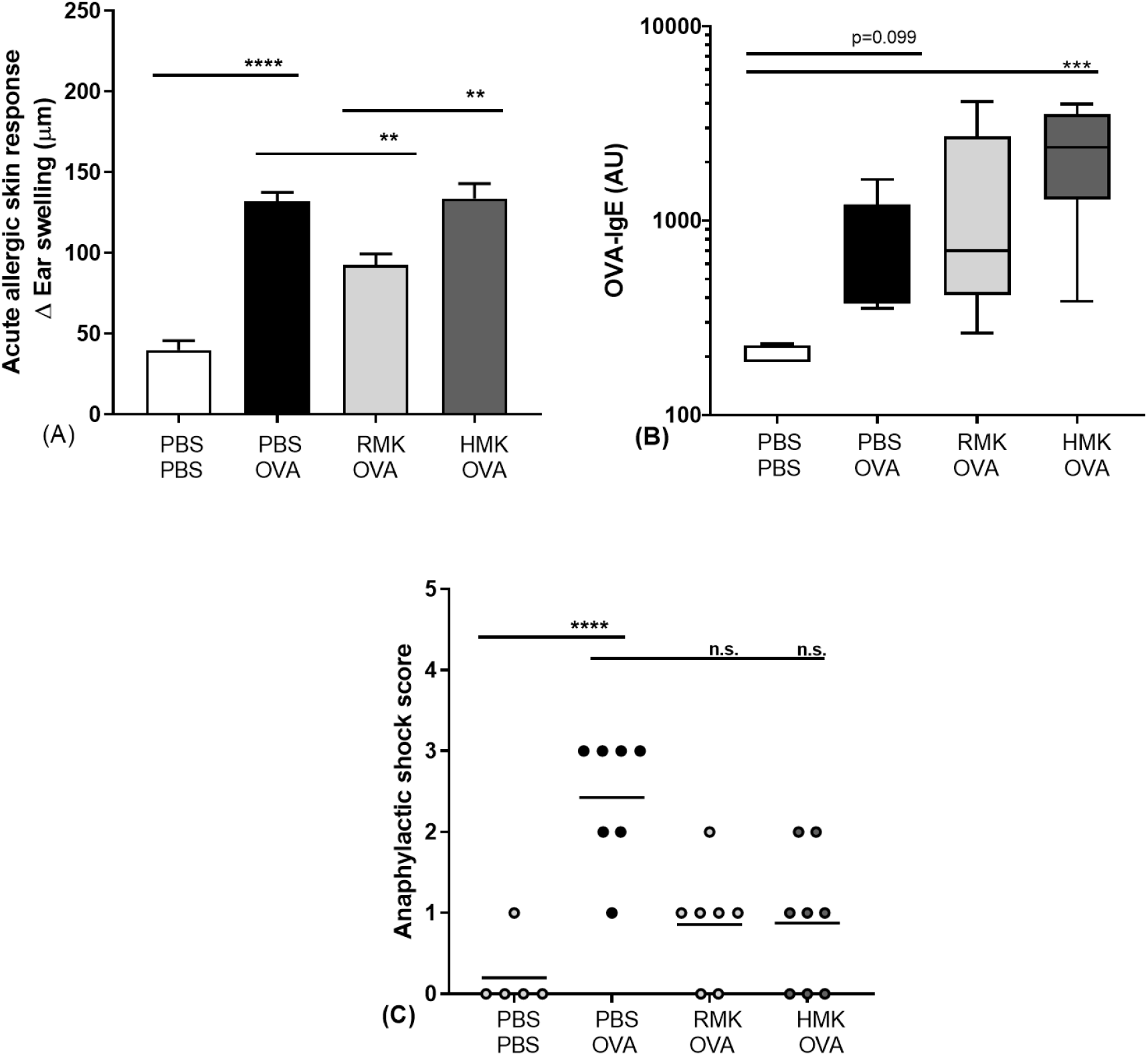
Reduced acute allergic skin response upon OVA challenge in mice treated with RMK. (A) The acute allergic skin response measured as Δ ear swelling 1 h after i.d. challenge, (B) OVA-specific IgE, (C) anaphylactic shock score. Data are presented as mean ± SEM for the acute allergic skin response, as box-and-whisker Tukey plot for OVA-IgE and as individual data points for anaphylactic shock scores, n = 6 in PBS group and n = 8 in all other groups. **P < 0.01, ***p<0.001, ****P < 0.0001, as analysed with one-way ANOVA followed by Bonferroni’s multiple comparisons test for pre-selected groups (A) or Kruskal-Wallis test for non-parametric data followed by Dunn’s multiple comparisons test for pre-selected groups (B, C). OVA = ovalbumin; RMK = raw milk kefir; HMK = heated milk kefir; i.d. = intradermal; n.s. = not significantly different.

#### Increased levels of OVA-specific IgE levels in mice treated with HMK

Allergic symptoms are predominantly mediated via allergen specific IgE. Therefore OVA-specific IgE and -IgG1 levels were measured in serum 16 hours after oral challenge. OVA-specific IgE levels were increased in PBS/OVA allergic mice compared to PBS/PBS sham sensitised mice (Fig 7B). Although not significantly different, low levels OVA-IgE were observed in the RMK/OVA mice and higher levels in the HMK/OVA mice. In contrast to RMK/OVA mice, OVA-IgE levels were significantly increased in HMK/OVA mice to PBS/PBS mice. No effect on OVA-IgG1 was observed in sensitised mice treated with RMK or HMK (data not shown).

#### Decreased activated Th1-cells and IFNg production after treatment with HMK

To further determine the local effects of the kefir treatments, T-cell subsets in MLN and cytokine production in spleen were studied. Percentages of activated Th2-, Th1-cells were not affected in PBS/OVA mice when compared to PBS/PBS mice (Fig 8A and 8C).

**Figure 8.**
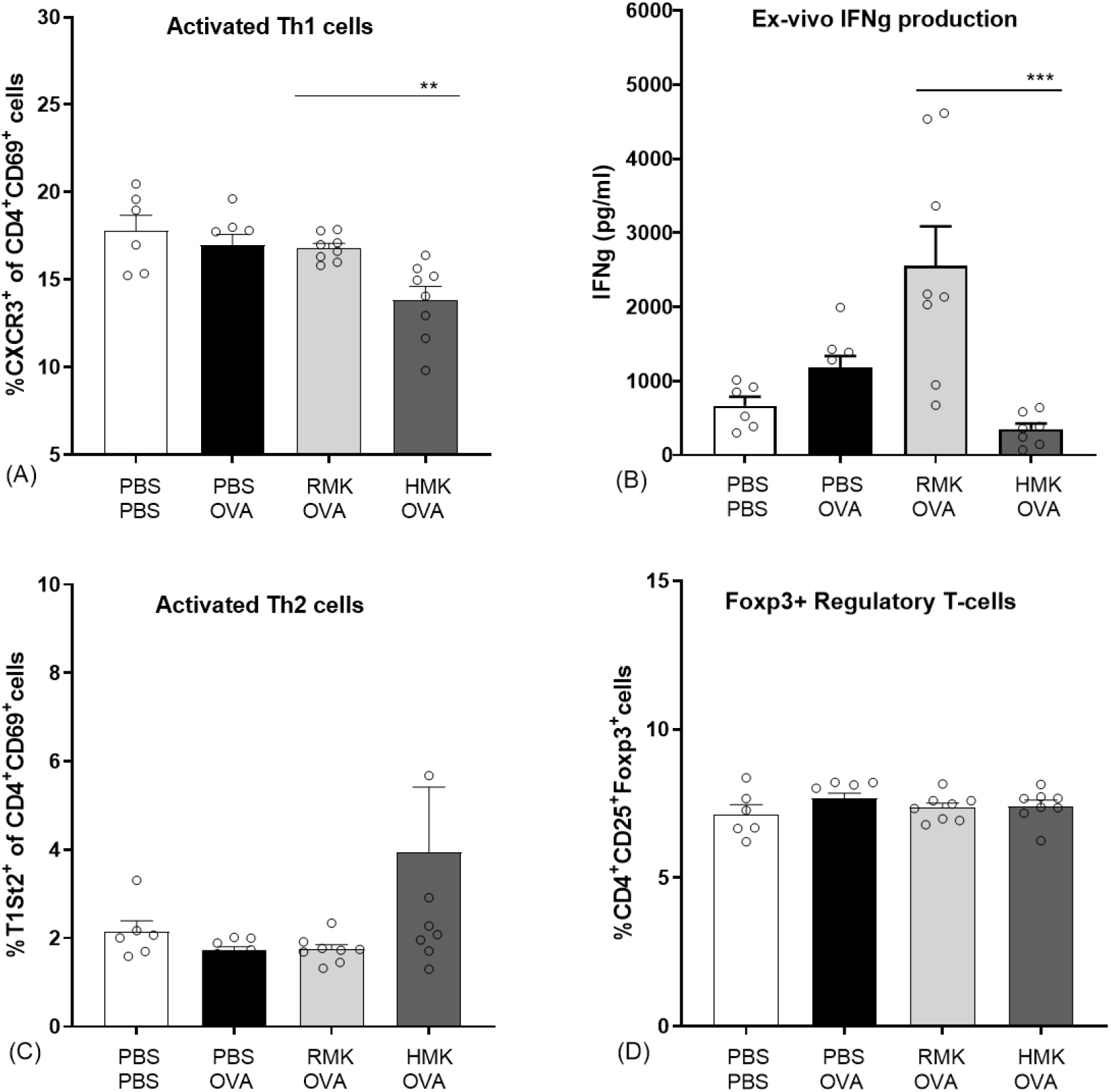
Decreased percentages of activated Th1-cells and IFNg production in mice treated with HMK. Percentages of (A) activated Th1 cells (CxCR3 of CD4+CD69+ cells) in MLN, (B) Concentration IFNg in ex vivo OVA stimulated splenocytes, (C) activated Th2 cells (T1ST2+ of CD4+ CD69+cells) and (D) Foxp3+ regulatory T-cells in MLN. Data are presented as mean ± SEM, n = 6 in PBS group and n = 8 in all other groups. **P < 0.01 as analysed with one-way ANOVA followed by Bonferroni’s multiple comparisons test for pre-selected groups. ***p<0.001 with Kruskal-Wallis test for non-parametric data followed by Dunn’s multiple comparisons test for pre-selected groups. OVA = ovalbumin; RMK = raw milk kefir; HMK = heated milk kefir; MLN = mesenteric lymph nodes.

Interestingly, HMK/OVA mice showed a reduction in activated Th1-cell percentages in MLN with no effects on the percentages of activated Th2-cells. The effect on activated Th1 cells was supported by a reduced IFNg production in splenocytes after *ex vivo* stimulation with OVA (Fig 8B). No effect was observed on Foxp3+ regulatory T-cells (Fig 8D) indicating that a shift in Th2/Th1 balance is underlying the protective effects.

## 4 Discussion

The safety of the RMK is based on two principles, a high hygienic standard during milking and processing as well as an immediate acidification of the raw milk. In several countries, legal raw milk has been produced under very strict hygienic conditions. The zoonotic risk is almost reduced till zero, and if any zoonotic bacteria were detected, there were no adverse health effects found by the authorities (38). As a result of the high hygienic standards applied at the dairy farm delivering the raw milk for this study, the bacterial contamination of the raw milk is kept around and below 10^3^ CFU/mL. Furthermore, there was a very short period between the end of milking and the start of culturing (less than two hours), leading to the rapid production of lactic acid, resulting in a lowering of the pH to 4.3, limiting the risk for microbial contamination and foodborne diseases (39,40), also in raw fermented milk (41). After 24h of fermentation with the added SC, kefir contains approximately 10^8^ bacteria and 10^5^ yeasts per mL. This agrees with previous reports, indicating ranges of lactic acid bacteria between 10^8^ and 10^9^ CFU/mL and yeast of 10^5^ and 10^6^ CFU/mL in final kefir products (42). With respect to storage, we observed a further increase of yeast counts over the period of three weeks, while the lactic acid bacteria reached their highest levels after one day of fermentation and numbers started to decline during storage, possibly due to further acidification. Indeed, other studies confirmed a slight decrease in lactic acid bacteria and increase of yeasts in kefir produced from a starter culture stored over a period of three weeks (43). Strikingly, the HMK showed higher levels of fungi than RMK after one day of fermentation. This could result from inactivation of the antifungal lactobacilli known to be present in RM (44).

Most of the detected microbial amplicons are identical to or are derived from the added SC and could be considered as safe or even beneficial for consumption. Although human infections with food derived yeasts harbouring virulence factors such as seen in some food derived from *Kluyveromyces marxianus* are potentially possible (45). *Streptococcus dysgalactiae* was detected in RM. It is most frequently encountered as a human commensal of the alimentary tract or genital tracts, and exposure to it could contribute to trained immunity (46).

Both RM and HM showed a very high richness of over one hundred microbial species. As these species have been identified based on their DNA in RM, this number may overrepresent the number of viable microbial species in the milk. Previous studies also led to the detection of hundreds of bacterial species in the RM microbiota. In agreement with the present study, members of the phyla Firmicutes and Proteobacteria account for most of these species (47). Their abundance depends on the specific environmental conditions. In our fresh raw milk microbiota, we found a rather even distribution of microbial species, in contrast to other studies where the milk-fermenting species *Streptococcus thermophilus* and *Lactococcus lactis* dominated the raw milk microbiota (48). The numbers of detected fungi were limited in our raw milk to about 30 ASVs, including the yeasts *Debaromyces, Mallasezia*, and *Candida* and the mould *Cladosporium*, all previously detected members of the raw milk mycobiota with a potential role in milk fermentation as some of them are efficient lactose degraders and show proteolytic and lipolytic activity (47). During the process of fermentation, the richness in microbial species drastically reduced. The most prominent members include the bacteria

*Streptococcus thermophilus, Lactococcus lactis, Lactococcus, Leuconostoc* and the yeast *Debaromyces* showing a perfect ASV-match to those of the microbial species present in the defined kefir starter culture. It should be noted that the kefir fermentation with a defined starter culture reported here is rather different from the kefir fermentation with natural grains, which includes a bacterial core community consisting of the kefir matrix former *Lactobacillus kefiranofaciens* and *Lactobacillus kefiri, Leuconostoc* sp., *Lactococcus lactis* and *Acetobacter* sp. (3). In the latter case a niche has been created by early fermenters by production of amino acids and lactate, which is converted to acetic acid by *Acetobacter* sp. Although the last process appears absent in our raw milk kefir production, as we did not identify any acetic acid bacteria, we did identify bacteria and yeasts that were not detectable or at low levels in early kefir fermentation and were more predominantly present after 6 to 12 hours of fermentation. These late fermenting microbial species included a specific variant of *L. lactis* and species of the fungi *Galactomyces* and *Pichia*. The former possibly uses galactose produced from lactose, while the latter species may be involved in proteolytic activity.

The production of kefir from raw milk produced in this study raises the question whether the raw milk microbiota participates in the kefir fermentation process. A comparative analysis of the microbiota from heated milk kefir indicates that several ASVs identified in raw milk kefir were absent in heated milk kefir. These ASVs match specific variants of the *L. lactis* and the fungus *Galactomyces geotrichum* and yeast *Pichia kudriavzevii*. It should be noted that the latter two species also have been identified in the raw milk which has been used for fermentation, whereas the variants of *L. lactis* were possibly below detection limit. Interestingly, co-cultures of *Galactomyces geotrichum* or *Pichia kudriavzevii* with *Lactobacillus* sp. showed a strong stimulating effect on the production of peptides (49). These peptides include bioactive anti-hypertensive peptides (50) with a strong activity against Angiotensin I-Converting Enzyme (ACE), a dipeptidyl carboxypeptidase that plays a role in the regulation of blood pressure (51). In addition, the presence of wild *L. lactis* strains in raw milk kefir could increase bioactive peptide levels in the final product (52).

The main result emerging from the peptide analysis is that upon fermentation, the number of unique peptides and the total peptide abundance increases by several orders of magnitude (Fig 5). This increase results from the proteolytic degradation by either endogenous or microbial proteolytic enzymes. The microbial proteases may originate both from the defined starter culture for both types of kefirs and the raw milk specifically for the RMK samples (7). Both before and after fermentation, most of the peptides were derived from the hydrolysis of caseins, especially β-casein (Table 1). For endogenous milk proteases like plasmin, it is known that caseins are their major target (53). Also, for lactic acid bacteria, it is known that their proteases preferentially hydrolyze caseins (54). Interestingly, additional bioactive peptides may be formed by the unique co-cultures of *Galactomyces geotrichum* or *Pichia kudriavzevii* with *Lactobacillus* sp. originating from raw milk, as discussed above (49). Many peptides present in milk were not identified after fermentation to kefir (Fig 5A), as 202 peptides were only found in the milk samples. The most probable explanation would be that these peptides are further degraded upon fermentation, leading to the formation of shorter peptides or free amino acids. This is underpinned by the peptide length data in Fig 5A, which shows that these 202 peptides only occurring in unfermented milk were on average longer than the peptides occurring only in kefir, or in both kefir and milk. When looking at the amino acids in the P1 and P1’ position of the peptides (supplemental file 5), there is a difference in cleavage specificity between RMK and HMK. However, as the proteases produced by the unique cultures in RMK are not characterized, it is not possible to determine whether there is a direct relation.

A comparative analysis of RMK and HMK showed that the number of unique peptides was higher in RMK than in HMK (172 *vs* 125; Fig 5A). This may be explained by the larger diversity in the microbiota of RMK (Fig 4), which could have led to proteolysis by a larger set of different proteases with different cleavage site specificities. Especially the presence of additional *Lactococcus lactis* or fungal strains in RMK may play a role, as it is known that these species produce a wide range of proteases and peptidases that will cleave casein at many different cleavage sites, leading amongst others to the formation of so-called peptide ladders (55), which were also found in our peptidomics dataset.

As studies on digestion of whey proteins have shown that denatured whey proteins are generally digested faster (56), we expected the HMK to contain a larger amount of whey protein-derived peptides. However, our data (Table 1 and Fig 6) do not confirm this hypothesis. As the sample underwent a short boiling step, which is a rather intense heat treatment, more extensive whey protein aggregation than during low pasteurisation may have occurred. It has previously been shown that the effect of whey proteins digestion depends on the specific aggregation structure ((57) and that intense heating may reduce whey protein proteolysis during fermentation (58). The specific, rather intense, heat treatment used in this study, may thus be the reason why no increase in whey protein-derived peptides was found in the HMK sample.

Nevertheless, although HMK showed to have less unique peptides than RMK, the total peptide abundance was higher. This is in line with the finding that proteolysis in HMK is carried out by a less diverse set of proteases, with less variation in cleavage specificity, leading to a less diverse peptide profile.

A search for known bioactivities showed that HMK has a higher bioactive potential than RMK when peptide abundances are considered (supplemental file 4). Nevertheless, for peptides that were identified only in HMK or RMK, most bioactive sequences were found in RMK. Although this could indicate a difference in the effect that HMK and RMK can have on health, caution is needed in interpretation since the extent of bioactivity is different for each peptide. Therefore, it is not possible to speculate on the role of specific peptides in an allergy-modulating effect of RMK.

In this study we show for the first time the protective effect of kefir on the food allergic response towards a food allergen. The outcomes are in accordance with the experiences in adults, consuming raw milk kefir. In a retrospective study, people experienced an improved score for their health, immunity, bowel, and mood after regular consumption of raw milk kefir, and health improvements were largest among people mentioning a poor health (59). In the underlying pre-clinical study, beneficial anti-allergic effects are only observed in kefir produced from raw unprocessed milk. A reduced acute allergic skin response is observed in RMK, but not in the kefir prepared from heated milk (HMK). The acute allergic skin response resembles the skin prick test in humans as used to identify sensitisation to specific allergens. Previous studies showed health benefits in relation to reduced inflammation and airway hyperresponsiveness in murine allergic asthma models (13,17). In those studies, kefir from pasteurised milk was used. The current study shows that only RMK possesses the putative capacity to redirect the food allergic response to the unrelated food allergen ovalbumin. Allergen-specific IgE is one of the key biomarkers responsible for the induction of allergic symptoms. Specific IgE is produced after sensitisation, the first encounter to the food allergen. Then IgE specifically binds to a high-affinity Fc receptor on the surface of mast cells or basophils that binds to allergen epitopes and triggers the release of inflammatory mediators and as a result the induction of food allergic symptoms. Although the effects of milk fermentation have been studied (60–62), only one study describes kefir related effects on food sensitisation. Hong et al showed suppression of IgE production and modulation of the T-cell compartment in LAB, *Lactobacillus kefiranofaciens* M1 treated mice (63). However, they did not investigated kefir as a matrix and measurement of clinically related symptoms were not included in the study. Although no difference was observed in OVA-IgE between RMK and HMK in the current study, higher OVA-specific IgE levels in HMK treated allergic mice compared to control mice might underly the protective effect of RMK. One of the main mechanisms leading to a heightened IgE response in food allergy is an imbalance in the Th1/Th2 cell (64–66). The allergy protective effect of RMK was related to a changed Th1/Th2 ratio as shown as an increase in activated Th1-cell percentages in mesenteric lymph nodes and splenic IFNg production in RMK treated mice (Fig 8). The change in Th1/Th2 ratio may be in part due to the increased number of regulatory T-cells. However, regulatory T-cell percentages are not changed in the current study.

## 5 Conclusion

The raw milk kefir made in this study was considered safe. Even though the microbial composition during fermentation changed towards the defined culture in kefir made from raw or heated milk, only in raw milk kefir specific microbes were also growing during fermentation. Their proteolytic activity may be responsible for the identified wider range of peptides in raw milk kefir. In line with these results, raw milk kefir reduced the acute allergic skin response and modulated T-cell responses in a murine food allergy model.

## Supplemental files

Supplemental file 1. Overview of the sample descriptions, cycle threshold values, and amplicon sequence variants.

Supplemental file 2. Fractions of microbial species in raw milk and heat-treated milk kefir.

Supplemental file 3. Precursor proteins of kefir peptides with the total peptide intensity (log10) in raw milk (RM), heated milk (HM), raw milk kefir (RMK), and heated milk kefir (HMK). Shown are proteins with at least 5 identified peptides in one of the 4 samples, arranged on the intensity in RM. Colour gradient represents high intensity (green) to low intensity (red).

Supplemental file 4. Overview of bioactivities found with the Milk Bioactive Peptide Database (MBPDB), ^(35)^ that match with identified peptide sequences.

Supplemental file 5. Relative intensities of amino acids in P1 and P1’ position for all peptides identified in (A) heated milk kefir (HMK) and (B) raw milk kefir (RMK). Relative intensities of amino acids in P1 and P1’ position for uniquely identified peptides in (C) HMK and (D) RMK.

## Author’s contributions

TB brought the research partners together. TB, BvE, KH and RK designed the research. TB, BvE, ZZ, PD, LvO, SB and MD conducted the research. BvE, ZZ, LvO and PD analysed the data. TB, BvE, KH, PD and RK wrote the paper. JG supervised the content. All authors read and approved the final manuscript.

## Acknowledgments

Software AG Stiftung, Darmstadt, Germany for funding the murine allergy study. Raw Milk Company, De Lutte, The Netherlands for providing the milk and kefir samples in this study as well as funding for the microbial enumeration experiments. Funding for DNA sequencing was obtained by Remco Kort from his NPN James Lind Award 2018.

## Abbreviations

ASV: Amplicon Sequence Variant
cat: Concatenated
CFU: Colony Forming Unit
Ct: Cycle threshold
SC: Defined Starter Culture eXact® 2 kefir, Hansen
EV: Extracellular Vesicles
ITS: Internal Transcribed Spacer
NCBI: National Centre for Biotechnology Information
PBS: Phosphate Buffered Saline solution
HM: Heated Milk (shortly boiled)
HMK: Heated Milk Kefir based on SC
qPCR: Quantitative real-time polymerase chain reaction
RA: Relative Abundance
RM: Raw Milk
RMK: Raw Milk Kefir based on SC

## References

1. Tamang JP, Cotter PD, Endo A, Han NS, Kort R, Liu SQ, et al. Fermented foods in a global age: East meets West. Compr Rev Food Sci Food Saf. 2020 Jan 1;19(1):184–217.

2. Gao W, Zhang L, Feng Z, Liu H, Shigwedha N, Han X, et al. Microbial diversity and stability during primary cultivation and subcultivation processes of Tibetan kefir. Int J Food Sci Technol. 2015 Jun 1;50(6):1468–76.

3. Blasche S, Kim Y, Mars RAT, Machado D, Maansson M, Kafkia E, et al. Metabolic cooperation and spatiotemporal niche partitioning in a kefir microbial community. Nat Microbiol. 2021 Feb 1;6(2):196–208.

4. Kiousi DE, Chorianopoulos N, Tassou CC, Galanis A. The Clash of Microbiomes: From the Food Matrix to the Host Gut. Microorganisms. 2022 Jan 1;10(1).

5. Marco ML, Heeney D, Binda S, Cifelli CJ, Cotter PD, Foligné B, et al. Health benefits of fermented foods: microbiota and beyond. Vol. 44, Current Opinion in Biotechnology. Elsevier Ltd; 2017. p. 94–102.

6. Ebner J, Aşçi Arslan A, Fedorova M, Hoffmann R, Küçükçetin A, Pischetsrieder M. Peptide profiling of bovine kefir reveals 236 unique peptides released from caseins during its production by starter culture or kefir grains. J Proteomics. 2015 Mar 8;117:41–57.

7. Dallas DC, Citerne F, Tian T, Silva VLM, Kalanetra KM, Frese SA, et al. Peptidomic analysis reveals proteolytic activity of kefir microorganisms on bovine milk proteins. Food Chem. 2016 Apr 15;197:273–84.

8. Bourrie BCT, Willing BP, Cotter PD. The microbiota and health promoting characteristics of the fermented beverage kefir. Vol. 7, Frontiers in Microbiology. Frontiers Media S.A.; 2016.

9. Kim DH, Jeong D, Kim H, Seo KH. Modern perspectives on the health benefits of kefir in next generation sequencing era: Improvement of the host gut microbiota. Vol. 59, Critical Reviews in Food Science and Nutrition. Taylor and Francis Inc.; 2019. p. 1782–93.

10. Slattery C, Cotter PD, O’Toole PW. Analysis of health benefits conferred by Lactobacillus species from kefir. Vol. 11, Nutrients. MDPI AG; 2019.

11. Farag MA, Jomaa SA, El-wahed AA, El-seedi HR. The many faces of kefir fermented dairy products: Quality characteristics, flavour chemistry, nutritional value, health benefits, and safety. Vol. 12, Nutrients. MDPI AG; 2020.

12. Azizi NF, Kumar MR, Yeap SK, Abdullah JO, Khalid M, Omar AR, et al. Kefir and its biological activities. Vol. 10, Foods. MDPI AG; 2021.

13. Lee MY, Ahn KS, Kwon OK, Kim MJ, Kim MK, Lee IY, et al. Anti-inflammatory and anti-allergic effects of kefir in a mouse asthma model. Immunobiology. 2007 Oct 15;212(8):647–54.

14. Tanaka M, Nakayama J. Development of the gut microbiota in infancy and its impact on health in later life. Allergology International. 2017 Oct;66(4):515–22.

15. Adiloğlu AK, Gönülateş N, Ìşler M, Şenol A. The Effect of Kefir Consumption on Human Immune System: A Cytokine Study. Mikrobiyol Bul. 2013 Apr 26;47(2):273–81.

16. Thoreux K, Schmucker DL. Kefir Milk Enhances Intestinal Immunity in Young but Not Old Rats. J Nutr. 2001 Apr 1;131(3):807–12.

17. Mendes E, Casaro MB, Fukumori C, Ribeiro WR, dos Santos AL, Sartorelli P, et al. Preventive oral kefir supplementation protects mice from ovariectomy-induced exacerbated allergic airway inflammation. Benef Microbes. 2021 Apr 12;12(2):187–97.

18. Brick T, Ege M, Boeren S, Böck A, von Mutius E, Vervoort J, et al. Effect of processing intensity on immunologically active bovine milk serum proteins. Nutrients. 2017 Sep 1;9(9):1–14.

19. Kontopodi E, Boeren S, Stahl B, van Goudoever JB, van Elburg RM, Hettinga K. High-Temperature Short-Time Preserves Human Milk’s Bioactive Proteins and Their Function Better Than Pasteurization Techniques With Long Processing Times. Front Pediatr. 2022 Jan 20;9.

20. Michalski MC, Januel C. Does homogenization affect the human health properties of cow’s milk? Trends Food Sci Technol. 2006 Aug;17(8):423–37.

21. Braun-Fahrländer C, von Mutius E. Can farm milk consumption prevent allergic diseases? Clinical and Experimental Allergy. 2011 Jan;41(1):29–35.

22. Loss G, Apprich S, Waser M, Kneifel W, Genuneit J, Büchele G, et al. The protective effect of farm milk consumption on childhood asthma and atopy: The GABRIELA study. Journal of Allergy and Clinical Immunology. 2011 Oct 1;128(4):766–73.

23. Abbring S, Ryan JT, Diks MAP, Hols G, Garssen J, van Esch BCAM. Suppression of food allergic symptoms by raw Cow’s milk in mice is retained after skimming but abolished after heating the milk—A promising contribution of alkaline phosphatase. Nutrients. 2019 Jul 1;11(7).

24. Abbring S, Xiong L, Diks MAP, Baars T, Garssen J, Hettinga K, et al. Loss of allergy-protective capacity of raw cow’s milk after heat treatment coincides with loss of immunologically active whey proteins. Food Funct. 2020 Jun 1;11(6):4982–93.

25. Abbring S, Kusche D, Roos TC, Diks MAP, Hols G, Garssen J, et al. Milk processing increases the allergenicity of cow’s milk—Preclinical evidence supported by a human proof-of-concept provocation pilot. Clinical and Experimental Allergy. 2019 Jul 1;49(7):1013–25.

26. Bolyen E, Rideout JR, Dillon MR, Bokulich NA, Abnet CC, Al-Ghalith GA, et al. Reproducible, interactive, scalable and extensible microbiome data science using QIIME 2. Nat Biotechnol. 2019 Aug 24;37(8):852–7.

27. Callahan BJ, McMurdie PJ, Rosen MJ, Han AW, Johnson AJA, Holmes SP. DADA2: High-resolution sample inference from Illumina amplicon data. Nat Methods. 2016 Jul 23;13(7):581–3.

28. Quast C, Pruesse E, Yilmaz P, Gerken J, Schweer T, Yarza P, et al. The SILVA ribosomal RNA gene database project: improved data processing and web-based tools. Nucleic Acids Res. 2012 Nov 27;41(D1):D590–6.

29. Nilsson RH, Larsson KH, Taylor AFS, Bengtsson-Palme J, Jeppesen TS, Schigel D, et al. The UNITE database for molecular identification of fungi: handling dark taxa and parallel taxonomic classifications. Nucleic Acids Res. 2019 Jan 8;47(D1):D259–64.

30. Lu J, Boeren S, de Vries SC, van Valenberg HJF, Vervoort J, Hettinga K. Filter-aided sample preparation with dimethyl labeling to identify and quantify milk fat globule membrane proteins. J Proteomics. 2011 Dec;75(1):34–43.

31. Dingess KA, de Waard M, Boeren S, Vervoort J, Lambers TT, van Goudoever JB, et al. Human milk peptides differentiate between the preterm and term infant and across varying lactational stages. Food Funct. 2017;8(10):3769–82.

32. Liu Y, de Groot A, Boeren S, Abee T, Smid EJ. Lactococcus lactis Mutants Obtained From Laboratory Evolution Showed Elevated Vitamin K2 Content and Enhanced Resistance to Oxidative Stress. Front Microbiol. 2021 Oct 14;12.

33. Cox J, Mann M. MaxQuant enables high peptide identification rates, individualized p.p.b.-range mass accuracies and proteome-wide protein quantification. Nat Biotechnol. 2008 Dec 30;26(12):1367–72.

34. Bateman A, Martin MJ, Orchard S, Magrane M, Agivetova R, Ahmad S, et al. UniProt: the universal protein knowledgebase in 2021. Nucleic Acids Res. 2021 Jan 8;49(D1):D480–9.

35. Nielsen SD, Beverly RL, Qu Y, Dallas DC. Milk bioactive peptide database: A comprehensive database of milk protein-derived bioactive peptides and novel visualization. Food Chem. 2017 Oct;232:673–82.

36. Wagih O. ggseqlogo: a versatile R package for drawing sequence logos. Bioinformatics. 2017 Nov 15;33(22):3645–7.

37. Li XM, Schofield BH, Huang CK, Kleiner GI, Sampson HA. A murine model of IgE-mediated cow’s milk hypersensitivity. J Allergy Clin Immunol. 1999 Feb;103(2 Pt 1):206–14.

38. Berge AC, Baars T. Raw milk producers with high levels of hygiene and safety. Epidemiol Infect. 2020;

39. O’Sullivan L, Ross RP, Hill C. Potential of bacteriocin-producing lactic acid bacteria for improvements in food safety and quality. Biochimie. 2002 May;84(5–6):593–604.

40. Nout MJR. Fermented foods and food safety. Food Research International. 1994 Jan;27(3):291–8.

41. Schoustra S, van der Zon C, Groenenboom A, Moonga HB, Shindano J, Smid EJ, et al. Microbiological safety of traditionally processed fermented foods based on raw milk, the case of Mabisi from Zambia. LWT. 2022 Sep;113997.

42. Hecer C, Ulusoy B, Kaynarca D. Effect of different fermentation conditions on composition of kefir microbiota. Vol. 26, International Food Research Journal. 2019.

43. Kök-Taş T, Seydim AC, Özer B, Guzel-Seydim ZB. Effects of different fermentation parameters on quality characteristics of kefir. J Dairy Sci. 2013 Feb;96(2):780–9.

44. Delavenne E, Mounier J, Déniel F, Barbier G, le Blay G. Biodiversity of antifungal lactic acid bacteria isolated from raw milk samples from cow, ewe and goat over one-year period. Int J Food Microbiol. 2012 Apr;155(3):185–90.

45. Peréz-Través L, de Llanos R, Flockhart A, García-Domingo L, Groenewald M, Pérez-Torrado R, et al. Virulence related traits in yeast species associated with food; Debaryomyces hansenii, Kluyveromyces marxianus, and Wickerhamomyces anomalus. Food Control. 2021 Jun;124:107901.

46. Netea MG, Domínguez-Andrés J, Barreiro LB, Chavakis T, Divangahi M, Fuchs E, et al. Defining trained immunity and its role in health and disease. Vol. 20, Nature Reviews Immunology. Nature Research; 2020. p. 375–88.

47. Quigley L, O’Sullivan O, Stanton C, Beresford TP, Ross RP, Fitzgerald GF, et al. The complex microbiota of raw milk. FEMS Microbiol Rev. 2013 Sep;37(5):664–98.

48. Masoud W, Vogensen FK, Lillevang S, Abu Al-Soud W, Sørensen SJ, Jakobsen M. The fate of indigenous microbiota, starter cultures, Escherichia coli, Listeria innocua and Staphylococcus aureus in Danish raw milk and cheeses determined by pyrosequencing and quantitative real time (qRT)-PCR. Int J Food Microbiol. 2012 Feb;153(1–2):192–202.

49. Chaves-López C, Serio A, Paparella A, Martuscelli M, Corsetti A, Tofalo R, et al. Impact of microbial cultures on proteolysis and release of bioactive peptides in fermented milk. Food Microbiol. 2014 Sep;42:117–21.

50. Hernández-Ledesma B, Miralles B, Amigo L, Ramos M, Recio I. Identification of antioxidant and ACE-inhibitory peptides in fermented milk. J Sci Food Agric. 2005 Apr 30;85(6):1041–8.

51. Riordan JF. Angiotensin-I-converting enzyme and its relatives [Internet]. Vol. 4, Genome Biology. 2003. Available from: http://genomebiology.com/2003/4/8/225

52. Rodríguez-Figueroa JC, González-Córdova AF, Torres-Llanez MJ, Garcia HS, Vallejo-Cordoba B. Novel angiotensin I-converting enzyme inhibitory peptides produced in fermented milk by specific wild Lactococcus lactis strains. J Dairy Sci. 2012 Oct;95(10):5536–43.

53. Ismail B, Nielsen SS. Invited review: Plasmin protease in milk: Current knowledge and relevance to dairy industry. Vol. 93, Journal of Dairy Science. 2010. p. 4999–5009.

54. Law J, Haandrikman A. Proteolytic enzymes of lactic acid bacteria. Int Dairy J. 1997 Jan;7(1):1–11.

55. Tjwan Tan PS, Poolman B, Konings WN. Proteolytic enzymes of Lactococcus lactis. Journal of Dairy Research. 1993 May 1;60(2):269–86.

56. Miralles B, del Barrio R, Cueva C, Recio I, Amigo L. Dynamic gastric digestion of a commercial whey protein concentrate†. J Sci Food Agric. 2018 Mar;98(5):1873–9.

57. Singh TK, Øiseth SK, Lundin L, Day L. Influence of heat and shear induced protein aggregation on the in vitro digestion rate of whey proteins. Food Funct. 2014;5(11):2686–98.

58. Guyomarc’h F. Formation of heat-induced protein aggregates in milk as a means to recover the whey protein fraction in cheese manufacture, and potential of heat-treating milk at alkaline pH values in order to keep its rennet coagulation properties. A review. Lait. 2006 Jan 1;86(1):1–20.

59. Baars T, Berge C, Garssen J, Verster J. The impact of raw fermented milk products on perceived health and mood among Dutch adults. Nutr Food Sci. 2019 Dec 12;49(6):1195–206.

60. Velez E, Novotny-Nuñez I, Correa S, Perdigón G, Maldonado-Galdeano C. Modulation of gut immune response by probiotic fermented milk consumption to control IgE in a respiratory allergy model. Benef Microbes. 2021 Apr 12;12(2):175–86.

61. Zhao L, Shi F, Xie Q, Zhang Y, Evivie SE, Li X, et al. Co-fermented cow milk protein by Lactobacillus helveticus KLDS 1.8701 and Lactobacillus plantarum KLDS 1.0386 attenuates its allergic immune response in Balb/c mice. J Dairy Sci. 2022 Sep;105(9):7190–202.

62. Pi X, Yang Y, Sun Y, Cui Q, Wan Y, Fu G, et al. Recent advances in alleviating food allergenicity through fermentation. Crit Rev Food Sci Nutr. 2022;62(26):7255–68.

63. Hong WS, Chen YP, Chen MJ. The antiallergic effect of kefir Lactobacilli. J Food Sci. 2010 Oct;75(8):H244–53.

64. Alashkar Alhamwe B, Meulenbroek LAPM, Veening-Griffioen DH, Wehkamp TMD, Alhamdan F, Miethe S, et al. Decreased Histone Acetylation Levels at Th1 and Regulatory Loci after Induction of Food Allergy. Nutrients. 2020 Oct 19;12(10).

65. Acevedo N, Alashkar Alhamwe B, Caraballo L, Ding M, Ferrante A, Garn H, et al. Perinatal and Early-Life Nutrition, Epigenetics, and Allergy. Nutrients. 2021 Feb 25;13(3):724.

66. Li QS, Wang YQ, Liang YR, Lu JL. The anti-allergic potential of tea: a review of its components, mechanisms and risks. Food Funct. 2021 Jan 7;12(1):57–69.

