## Supplemental file 3 for "Raw milk kefir: microbiota, bioactive peptides, and immune modulation"

*Precursor proteins of kefir peptides with the total peptide intensity (log10) in raw milk (RM), heated milk (HM), raw milk kefir (RMK), and heated milk kefir (HMK). Shown are proteins with at least 5 identified peptides in one of the 4 samples, arranged on the intensity in RM. Colour gradient represents high intensity (green) to low intensity (red).*

| Protein name | RM | HM | RMK | HMK |
| --- | --- | --- | --- | --- |
| α_S1_-casein | 10.4 | 10.5 | 11.6 | 11.6 |
| Glycosylation-dependent cell adhesion molecule 1 | 10.1 | 10.0 | 11.4 | 11.2 |
| α_S2_-casein | 10.1 | 10.0 | 11.1 | 11.2 |
| β-casein | 10.0 | 9.9 | 12.0 | 12.1 |
| Polymeric immunoglobulin receptor | 9.4 | 9.4 | 10.4 | 10.4 |
| Perilipin-2 | 8.8 | 8.8 | 8.7 | 8.2 |
| Butyrophilin subfamily 1 member A1 | 8.8 | 8.8 | 8.9 | 8.9 |
| Glycoprotein 2 | 8.7 | 8.7 | 6.7 | 8.4 |
| Sodium-dependent phosphate transport protein 2B | 8.6 | 8.6 | 9.0 | 9.0 |
| Lactoperoxidase | 8.6 | 8.6 | 7.5 | 8.2 |
| Kappa-casein | 8.4 | 9.3 | 11.6 | 11.9 |
| Osteopontin | 8.0 | 8.0 | 9.9 | 9.9 |
| β-lactoglobulin | 8.0 | 9.7 | 7.8 | 8.5 |
| β-2-microglobulin | NA | NA | 9.5 | 9.2 |
| Nucleobindin-1 | NA | NA | 8.6 | 8.2 |
