## Supplemental file 4 for "Raw milk kefir: microbiota, bioactive peptides, and immune modulation"

*Overview of bioactivities found with the Milk Bioactive Peptide Database (MBPDB)* *that match with identified peptide sequences* (Nielsen et al., 2017).

|  | Number of times bioactivities were found for peptide sequences identified: | | | |
| --- | --- | --- | --- | --- |
| Bioactivity | More abundant in HMK | More abundant in RMK | Uniquely in HMK | Uniquely in RMK |
| ACE-inhibitory | 328 | 255 | 40 | 63 |
| Antioxidant | 173 | 119 | 20 | 39 |
| Antimicrobial | 143 | 121 | 18 | 34 |
| DPP-IV inhibitory | 92 | 85 | 7 | 23 |
| Anti-inflammatory | 51 | 15 | 15 | 6 |
| Opioid | 48 | 27 | 2 | 9 |
| Immunomodulatory | 37 | 32 | 5 | 12 |
| Anticancer | 26 | 22 | 1 | 7 |
| Inhibition of cholesterol solubility | 20 | 14 | 2 | 4 |
| Prolyl endopeptidase-inhibitory | 20 | 19 | 0 | 6 |
| Osteoanabolic | 18 | 13 | 5 | 2 |
| Bradykinin-potentiating | 17 | 3 | 0 | 0 |
| Increase MUC4 expression | 12 | 4 | 1 | 1 |
| Growth-promoting | 11 | 4 | 2 | 2 |
| Anxiolytic | 10 | 10 | 0 | 4 |
| Cytomodulatory | 10 | 11 | 2 | 3 |
| Antithrombotic | 8 | 4 | 2 | 3 |
| Ameliorates insulin resistance | 6 | 1 | 2 | 0 |
| Antihypertensive | 6 | 3 | 0 | 1 |
| Increases jejunal mucus secretion | 6 | 8 | 0 | 3 |
| Increases MUC2 expression | 6 | 8 | 0 | 3 |
| Increases MUC3 expression | 6 | 8 | 0 | 3 |
| Increases MUC5a expression | 6 | 8 | 0 | 3 |
| Reduces pancreas MDA level | 6 | 8 | 0 | 3 |
| Satiety | 6 | 8 | 0 | 3 |
| Cathepsin B inhibitory | 5 | 7 | 0 | 3 |
| Antithrombin | 4 | 0 | 1 | 0 |
| Anti-apoptotic effect | 3 | 2 | 1 | 0 |
| Wound healing | 3 | 2 | 1 | 0 |
| Cytotoxic | 2 | 0 | 0 | 0 |
| Non-functional | 2 | 6 | 0 | 2 |
| Cancer | 1 | 3 | 0 | 1 |
| Improves learning and memory | 1 | 3 | 0 | 1 |
| Increases intestinal motility | 1 | 3 | 0 | 1 |
| Promote neurite outgrowth | 1 | 3 | 0 | 1 |
| Stimulates proliferation | 1 | 0 | 0 | 0 |
| Increase small intestinal goblet cell density | 0 | 1 | 0 | 0 |
| Promote calcium uptake | 0 | 2 | 0 | 1 |
| Protective effects in indomethacin-induced enteritis through preservation of goblet cells and improvement in wound healing | 0 | 1 | 0 | 0 |
