## Supplemental file 5 for "Raw milk kefir: microbiota, bioactive peptides, and immune modulation"

*Relative intensities of amino acids in P1 and P1’ position for all peptides identified in (A) heated milk kefir (HMK) and (B) raw milk kefir (RMK). Relative intensities of amino acids in P1 and P1’ position for uniquely identified peptides in (C) HMK and (D) RMK.*

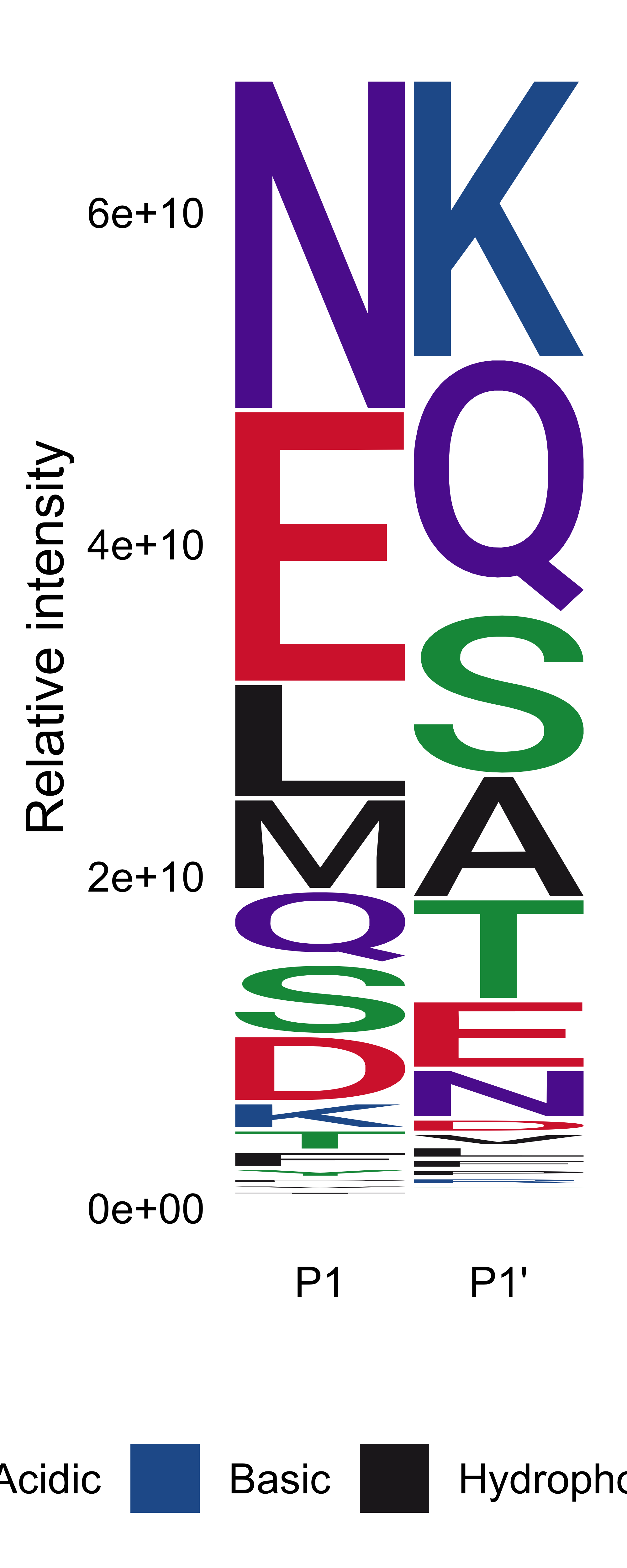

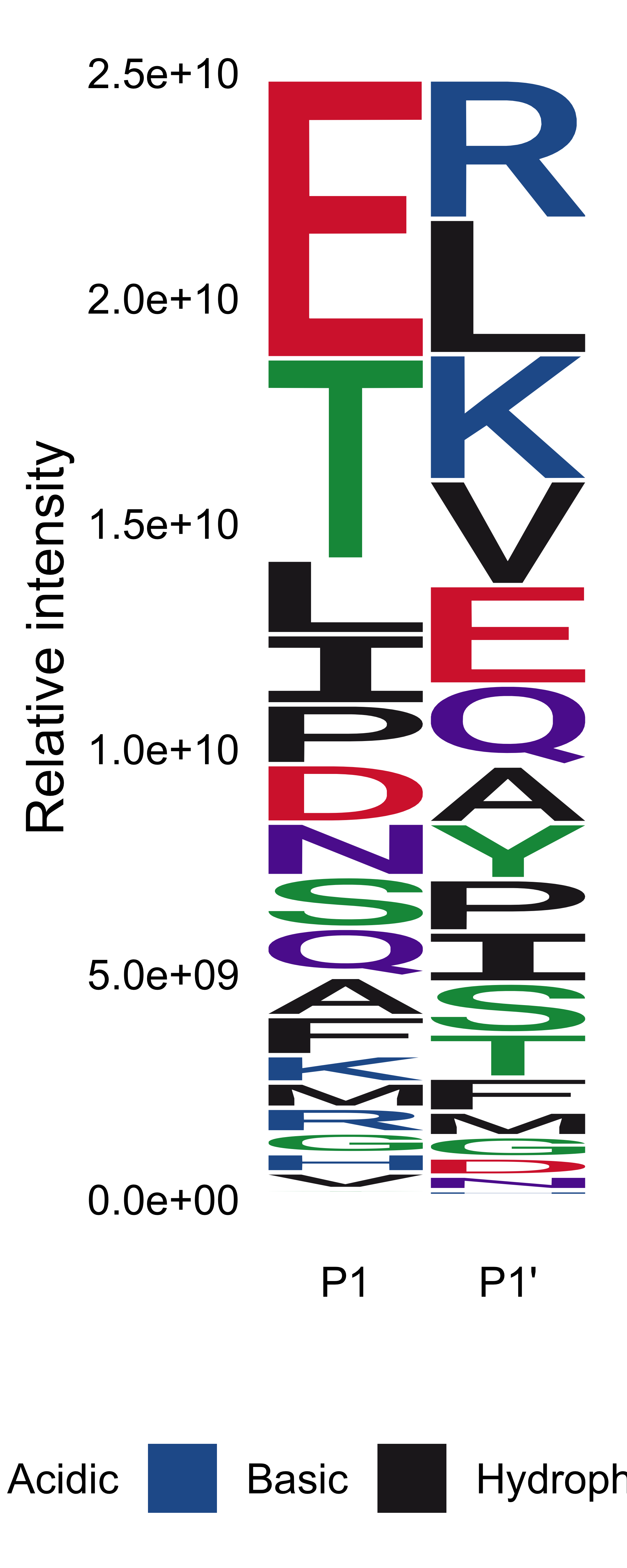

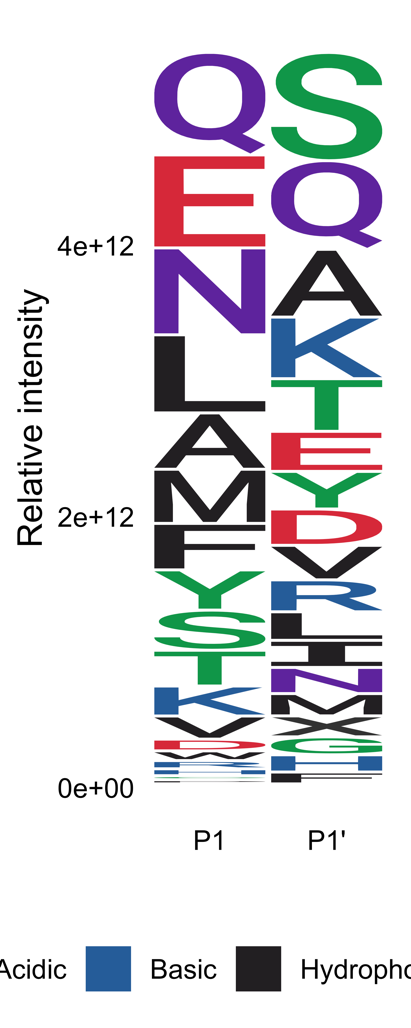

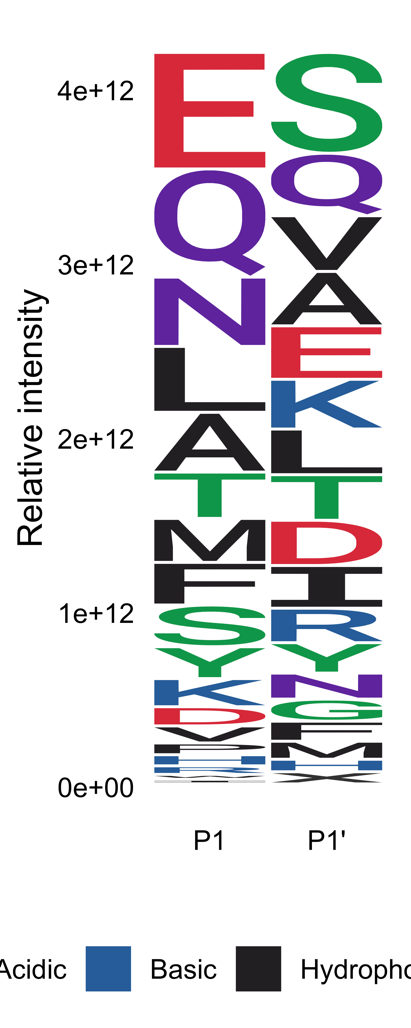

A

B

C

D

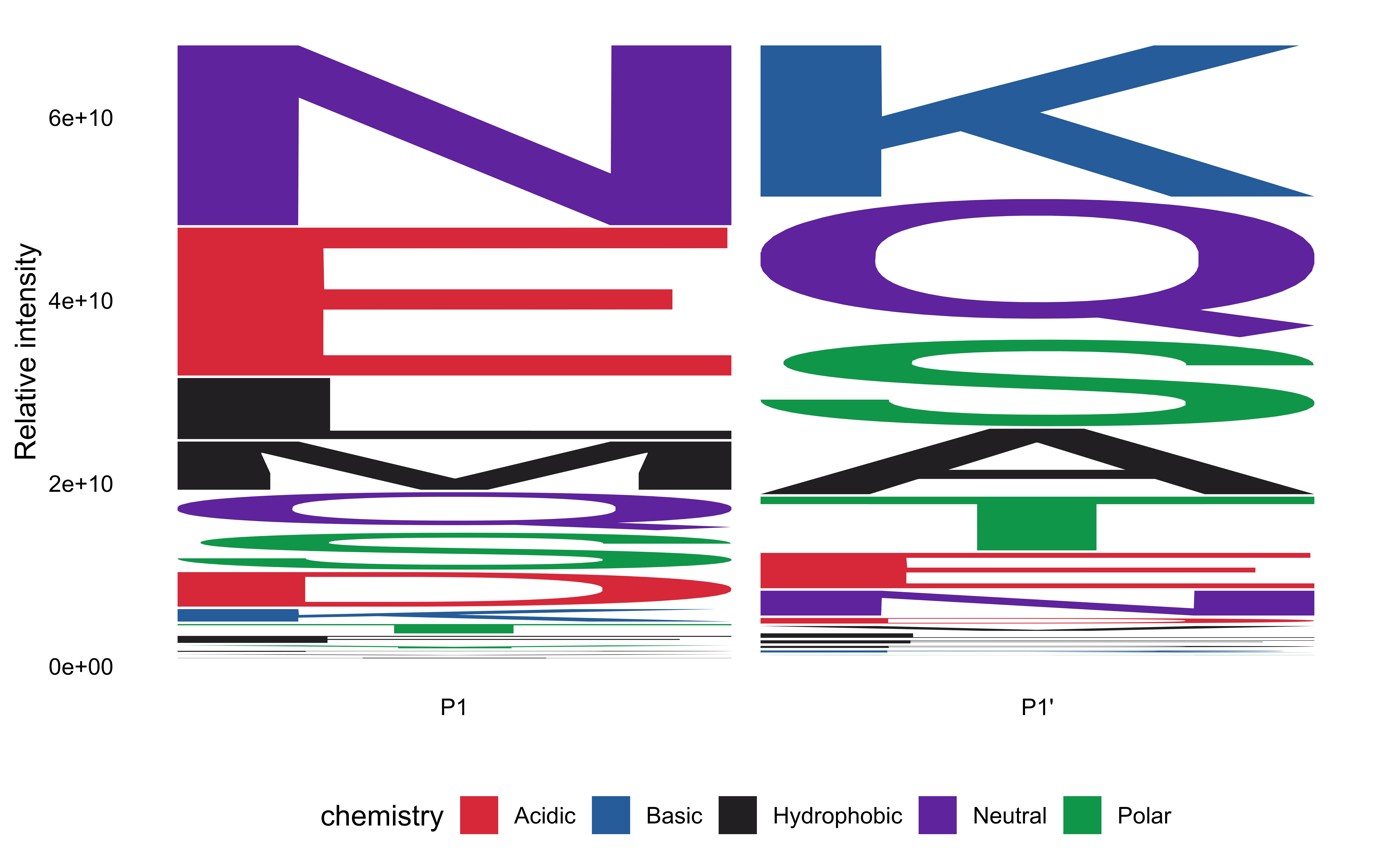
